# The unique ORF8 protein from SARS-CoV-2 binds to human dendritic cells and induces a hyper-inflammatory cytokine storm

**DOI:** 10.1101/2022.06.06.494969

**Authors:** Matthias Hamdorf, Thomas Imhof, Ben Bailey-Elkin, Janina Betz, Sebastian J Theobald, Alexander Simonis, Veronica Di Cristanziano, Lutz Gieselmann, Felix Dewald, Clara Lehmann, Max Augustin, Florian Klein, Miguel A Alejandre Alcazar, Robert Rongisch, Mario Fabri, Jan Rybniker, Heike Goebel, Jörg Stetefeld, Bent Brachvogel, Claus Cursiefen, Manuel Koch, Felix Bock

## Abstract

The novel coronavirus pandemic, whose first outbreak was reported in December 2019 in Wuhan, China (COVID-19), is caused by the <u>s</u>evere <u>a</u>cute respiratory <u>s</u>yndrome <u>c</u>oronavirus <u>2</u> (SARS-CoV-2). Tissue damage caused by the virus leads to a strong immune response and activation of antigen-presenting cells, which can elicit acute respiratory distress syndrome (ARDS) characterized by the rapid onset of widespread inflammation, the so-called cytokine storm. In many viral infections the recruitment of monocytes into the lung and their differentiation to dendritic cells (DCs) are seen as a response to the viral infection. DCs are critical players in the development of the acute lung inflammation that causes ARDS. Here we focus on the interaction of the ORF8 protein, a specific SARS-CoV-2 open reading frame protein, with dendritic cells (DCs). We show that ORF8 binds to dendritic cells, causes a pre-maturation of differentiating DCs, and induces the secretion of multiple pro-inflammatory cytokines by these cells. In addition, we identified dendritic cell-specific intercellular adhesion molecule-3-grabbing non-integrin (DC-SIGN) as a possible interaction partner of ORF8 on dendritic cells. Blockade of ORF8 signaling leads to reduced production of IL-1β, IL-6, IL-12p70, TNF-α, MCP-1 (CCL2), and IL-10 by dendritic cells. Analysis of patient sera with high anti-ORF8 antibody titers showed that there was nearly no neutralization of the ORF8 protein and its function. Therefore, a neutralizing antibody that has the capacity of blocking the cytokine and chemokine response mediated by ORF8 protein might be an essential and novel additional step in the therapy of severe SARS-CoV-2 cases.

## Introduction

The novel coronavirus pandemic, whose first outbreak was reported in December 2019 in Wuhan, China (COVID-19), is caused by the <u>s</u>evere <u>a</u>cute respiratory <u>s</u>yndrome <u>c</u>oronavirus <u>2</u> (SARS-CoV-2) (1–3). This virus has risen to a global pandemic with over ∼267.208 million infections and over ∼ 5.272 million death (John Hopkins University, 8^th^ December 2021). The SARS-CoV-2 is a positive-stranded enveloped RNA virus that belongs to the family of Coronaviridae, a virus family with a natural reservoir in bats that has 55% nucleotide similarity and 30% protein sequence similarity with SARS-CoV, the virus that caused the first documented previous outbreak of severe acute respiratory syndrome in 2002 (4). During acute infection, the cytokine release syndrome (CRS) seems to be responsible for the severe conditions and the development of SARS (5). The disease is divided into two phases. During the non-severe stage, the virus triggers the innate and adaptive immune response. The innate immunity is mainly mediated by phagocytic cells, i.e. professional antigen-presenting cells, including dendritic cells (DCs), macrophages, and granulocytes that are residential in the lung or infiltrate into the lung tissue after infection (5). Especially the recruitment of monocytes into the lung and their differentiation into inflammatory, CD1c+, monocyte derived DCs (moDCs) plays a crucial role in viral infection like influenza (6) as well as SARS-CoV-2 (7, 8). During this phase, the body begins to control the virus infection by an orchestrated T and B cell response and starts to eliminate the virus to suppress disease progression. However, if the body fails to develop a protective immune response during this early phase, the virus propagates, and massive destruction of the affected tissues occurs in the severe stage (9). The tissue damage caused by the virus leads to a strong response and activation of antigen-presenting cells. This second innate immune response can elicit acute respiratory distress syndrome (ARDS), which causes life-threatening respiratory disorders and is characterized by the rapid onset of widespread inflammation, the so-called cytokine storm (10). Dendritic cells are important players among other innate and adaptive immune cells in the development of the acute lung inflammation that causes ARDS (5, 11, 12) and a related cytokine storm. Therefore, these cells are center of this study.

Even if patients are not running into a severe acute respiratory syndrome with an uncontrolled inflammatory storm, SARS-CoV-2 infected patients suffer from a massive variety of long-term symptoms (13). The over-reactive immune system triggered by the infection leads to inflammation, and one particularly susceptible organ that is affected is the heart. One of the symptoms induced by inflammation is cardiomyopathy, in which the muscles of the heart become stretched, stiff, or thickened, affecting the heart’s ability to pump blood. These symptoms are occurring in patients even after discharge from the hospital (14). The study of Carfi, A. et al. with 143 people that survive the infection with COVID-19 report as the most common lasting effect fatigue (53%) and shortness of breath (43%) (15). In 88% of the patients, visible damages of the lung are still detectable even six weeks after being discharged from the hospital, and even after 12 weeks, the number is only decreasing to 56%. The symptoms that are related to this long-lasting damage may take a longer time to fade and may still trigger a chronic inflammation (16, 17). For the SARS-CoV-2 virus, the ongoing inflammation and the long-term clinical picture seems to differ from the two other known coronavirus infections in human, SARS-CoV and MERS-CoV (18). This leads to the question of how these viruses differ from each other.

The SARS-CoV-2 virus contains a unique new open reading frame 8 (*orf8*) gene that differs from the reading frames in SARS-CoV-1 and MERS-CoV (19). Several functions have been predicted for the ORF8 protein of SARS-CoV-2. In cell culture, ORF8 exogenous overexpression disrupts IFN-I signaling (20). It has been shown that ORF8 of SARS-CoV-2, but not ORF8 or ORF8a/b of SARS-CoV-1, is involved in the downregulation of MHC-I in cells (21).

A recent study focused on the secretion of the ORF8 protein from infected cells in vitro as well as secreted ORF8 protein in patients. Wang et al. showed that ORF8 is mostly a secretory protein. Additionally, the study showed that patients with a SARS-CoV-2 infection showed early seropositivity for anti-ORF8 IgM, IgG, and IgA and that ORF8 can be used as an early diagnostic marker (22). Since dendritic cells (DCs) and macrophages act as antigen-presenting cells (APCs), the infection of these cells by SARS-CoV-2 or the interaction with virus proteins may impair the adaptive immune responses against the virus and trigger the initial process of the cyto-/chemokine storm (23). The function and interaction of dendritic cells with virus proteins and their response are still poorly understood. Therefore, our study focus is on the interaction of the secreted ORF8 protein with dendritic cells and its contribution to the cytokine storm observed in COVID-19 patients (24).

## Results

### ORF8 specifically binds to monocytes and dendritic cells

ORF8 contains a short signal peptide sequence similar to the one present in the S-protein. Recently, it has been shown that ORF8 is secreted from infected cells, indicating the functionality of the signal peptide sequence (22). To confirm this result, we transfected HEK293 cells with the wild type ORF8 coding sequence, including the short 5’ untranslated region. After induction for 24 h with doxycycline, ORF8 was detectable in the supernatant by coomassie blue staining (Figure S2). Since CD14+ monocytes are able to recognize foreign proteins, we analyzed if the ORF8 protein can interact with this cell type. Therefore we used a recombinant ORF8 protein (Figure S1 a, b) labeled with Atto488. We could show that within human peripheral blood mononuclear cells (PBMCs), mainly CD14+ monocytes bound ORF8 (Figure 1B, n=3 and Figure S3 a). To exclude that this binding is a random protein binding we used labled BSA as a control. The binding of ORF8 to monocytes and related DCs is statistically much higher than to a random protein (Figure S3 a, b). As monocytes are the origin of the antigen-presenting dendritic cells (Figure 1 a), we followed up the binding of ORF8 to immature and mature dendritic cells. Both populations (n=3 donors for each experiment) have the ability to interact with the ORF8 protein, as shown in Figure 1 c and Figure S3 c. Monocytes, as well as dendritic cells, are known to bind and uptake antigens of a considerable variety of sources. To validate this specific binding of ORF8, we tested a commercially available rabbit anti-ORF8 polyclonal antibody for blocking. As shown, this polyclonal antibody led to reduced binding of ORF8 to immature and mature dendritic cells (Figure 1 d), confirming the specificity of this interaction. The corresponding isotype controle was not able to reduce ORF8 binding (Figure S3 d) and the BSA-Atto488 protein showed only a very week interaction with monocytes (Figure S3 a) and related dendritic cells (Figure S3 b).

**Figure 1:**
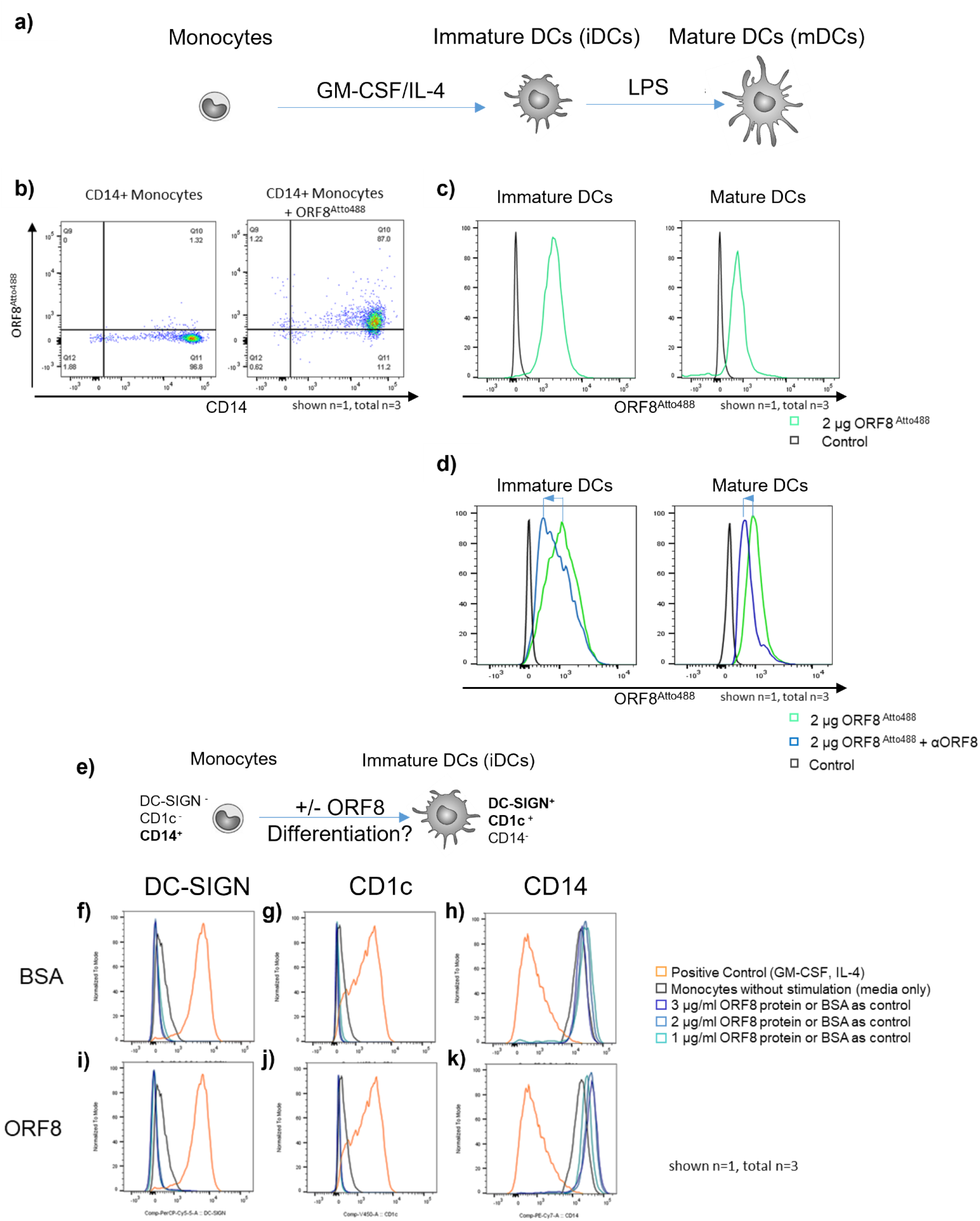
ORF8 binds specifically to monocytes and DCs and is lacking the property to induce DC differentiation. a) Experimental setup of differentiation and maturation of dendritic cells from monocytes; b) flow cytometric analysis of CD14+ monocytes within PBMCs on the binding of ORF8-Atto488; negative control: unstained cells; c) flow cytometric analysis of binding of ORF8-Atto488 to immature and mature DCs; d) flow cytometry analysis of the effect of a polyclonal anti-ORF8 antibody (rabbit) on the binding of ORF8 to immature and mature dendritic cells (blue arrows indicates the shift of peak); e) Experimental setup: CD14+ monocytes were treated with ORF8 and analyzed for markers of immature monocyte-derived dendritic cells (immature MoDCs); f-k) Flow cytometry analysis of the expression of DC-SIGN (f, i), CD1c (g, j), and CD14 (h, k) on monocytes after incubation with ORF8 (i-k) compared to the incubation with BSA (f-h); as positive control monocytes were differentiated to immature dendritic cells by GM-CSF and IL-4 (orange line); monocytes without any stimulation served as negative control (black line); representative flow cytometry histograms (n=3 donors)

### ORF8 overdrives DC differentiation into pre-maturity

Dendritic cells are an important modulator of the immune response and may play an important role in the clearance of SARS-CoV-2 infections. As monocytes are recruited into the lung during infection and differentiate into inflammatory, monocyte derived DCs, we aimed to define the function of ORF8 as an immune modulator and virulence factor in the following part. In dendritic cell biology, antigens and especially superantigens from some viruses and bacteria can induce the differentiation of monocytes to dendritic cells (25, 26) (Figure 1 e). In our study, we found that ORF8 showed no influence on monocytes, similar to BSA, and is not acting as an inducer of differentiation (Figure 1 f-k; n=3). Monocytic precursor cells treated with BSA or ORF8 remained positive for the monocyte marker CD14 (Figure 1 h, k respectively) and negative for the dendritic cell markers DC-SIGN (Figure 1 f, i, respectively), and CD1c (Figure 1 g, j respectively).

Next, we analyzed if ORF8 can influence the differentiation process. Therefore, we differentiated monocytes to immature dendritic cells (immature MoDC) with GM-CSF and IL-4 in the presence or absence of ORF8 in different concentrations (Figure 2 a). We performed a dose escalation of ORF8 protein and looked for the upregulation of the maturation marker CD40 and CD80. A first plato phase was reached between 500 and 1000 ng/ml and had an additional small increase at 2000 ng/ml (Figure S4 c). Based on that we chose a concentration of 1000 ng/ml for all further experiments to elict a robust activation. ORF8 treated cells showed a significantly different stellate-like morphology which is comparable to mature dendritic cells (Figure S8 a, b). In line with the morphological changes, also the expression of the maturation markers MHCII, CD83, CD80, CD86, and CD40 were markedly upregulated. In contrast, the dendritic marker CD11c was slightly downregulated (Figure 2 b, total n=6). The surface markers of a conventional induced maturation are shown in Figure S8 c. Neither BSA, nor the polyclonal anti-ORF8 antibody alone, its isotype control or a human Fc-antibody fragment as controls were able to induce maturation (Figure S3 e).

**Figure 2:**
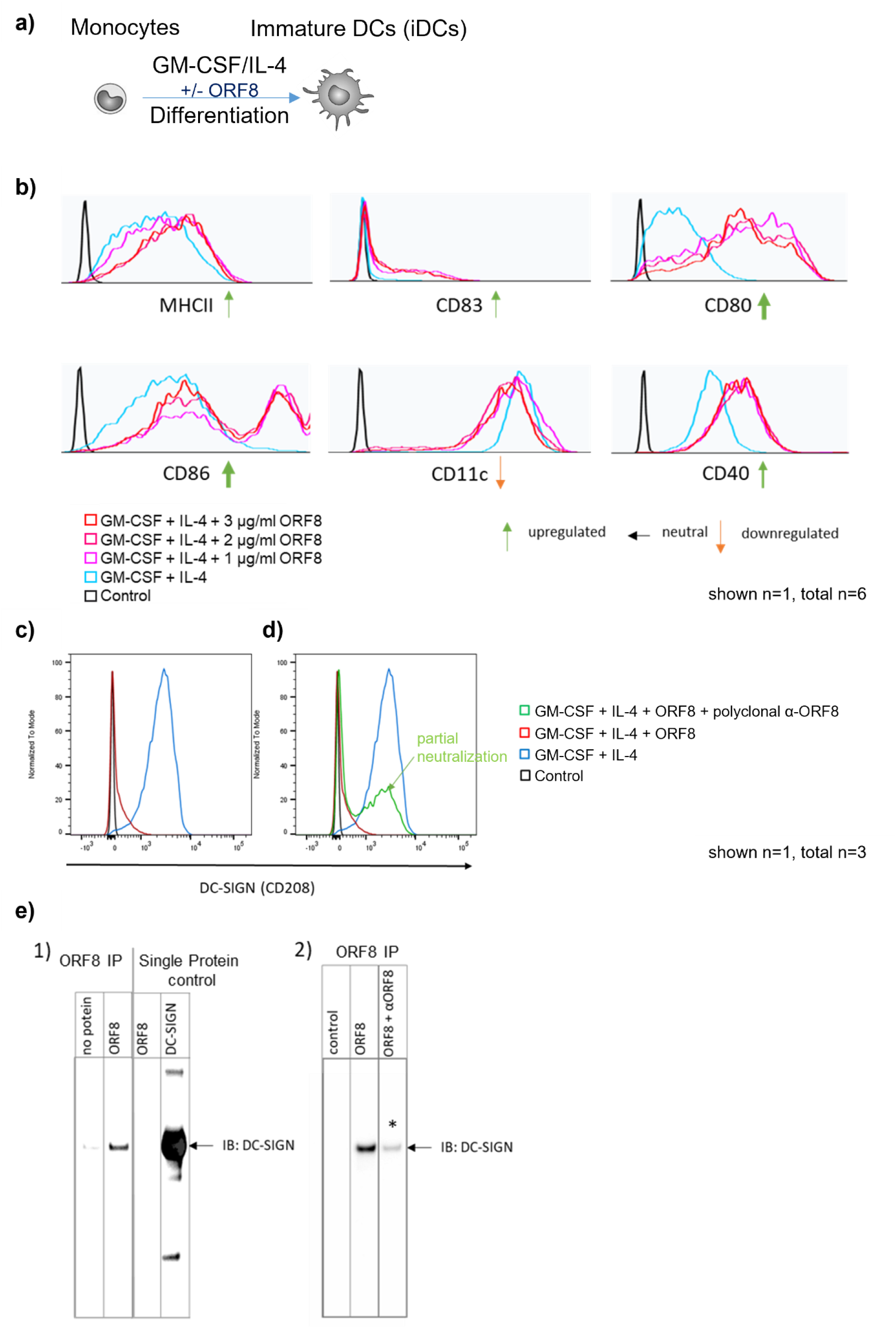
ORF8 overdrives DC differentiation into maturity/activation and downregulates DC-SIGN. a) Experimental set up: monocytes were differentiated into immature dendritic cells in presence or absence of ORF8; b) Flow cytometry analysis of the effect of ORF8 on the expression of MHCII, CD80, CD83, CD86, CD40 and CD11c during differentiation of monocytes into immature dendritic cells with IL-4 and GM-CSF compared to only IL-4+GM-CSF differentiated monocytes (blue line); control: monocytes without IL-4+GM-CSF treatment (black line); representative flow cytometry histograms (n=6 donors) c) Flow cytometry analysis of the effect of ORF8 on the expression of DC-SIGN during differentiation of monocytes into immature dendritic cells with IL-4 and GM-CSF (red line) compared to only IL-4+GM-CSF differentiated monocytes (blue line); control: monocytes without IL-4+GM-CSF treatment (black line); representative flow cytometry histograms (n=3 donors) d) Flow cytometry analysis of the effect of the blockade of ORF8 by an inhibitory anti-ORF8 antibody (α-ORF8-iAB) on the expression of DC-SIGN on differentiated dendritic cells compared to only IL-4+GM-CSF differentiated monocytes (blue line) and IL-4+GM-CSF+ORF8 differentiated monocytes (red line); control: monocytes without IL-4+GM-CSF treatment (black line; for isotype control see supplemental Figure S2d); representative flow cytometry histograms (n=3 donors); e) Immunoprecipitations (IP) of recombinant DC-SIG with 1) no protein control, recombinant ORF8, ORF8 + an inhibitory anti-ORF8 antibody (αORF8), 2) single recombinant proteins (single protein) ORF8 and recombinant human DC-SIGN (positive control) were immunoblotted (IB) with anti-DC-SIGN antibody; * indicates reduced signal intensity compared to ORF8 alone

In addition, we also analyzed the DC specific marker DC-SIGN in this experimental setup. Interestingly, the expression of this marker was almost completely downregulated (Figure 2 c; n=3).

To verify the specific effect of ORF8 on the expression of DC-SIGN we used a polyclonal anti-ORF8 antibody. Neutralizing ORF8 with this polyclonal antibody partially restored the expression of DC-SIGN. These results demonstrate that the effect is regulated by the ORF8 protein (Figure 2 d, n=3, labeled with the green arrow). Since DC-SIGN was shown to be important for the internalization of HIV (27), we next analyzed by immune precipitation if ORF8 interacts with DC-SIGN. We could show in a co-immunoprecipitation that ORF8 interacts with the DC-SIGN receptor (Figure 2 e, Blot 1). Using the polyclonal anti-ORF8 antibody again to neutralize ORF8, we could reduce the binding of DC-SIGN to ORF8 (Figure 2 e, Blot 2, marked with an asterisk), demonstrating that the anti-ORF8 antibody interferes with the DC-SIGN/ORF8 interaction side.

### ORF8 induces a unique pro-inflammatory cytokine secretion during DC differentiation

SARS-CoV-2 can induce a cytokine storm leading to ARDS (10). Therefore, we also analyzed the chemokine and cytokine expression profile of dendritic cells exposed to ORF8 during differentiation. Strikingly, we found a significant, strongly elevated secretion of IP-10, IL-1β, TNF-α, MCP-1, IL-6, IL-10, and IL12p70 (Figure 3 a) as well as an increased level of IFN-γ and IL-8 (Fig. S12 b) in comparison to untreated cells (0 µg/ml ORF8). This effect could be partly reversed by the simultaneous addition of a neutralizing polyclonal rabbit antibody against ORF8 during the differentiation, verifying the ORF8 specific induction of the cytokine storm (Figure S6). Parallel to the exclusion of contamination of the purified ORF8 protein with endotoxin (Figure S1 c), we could also show that the ORF8 mediated cytokine storm has a unique fingerprint compared to the cyto-/chemokine profile induced by LPS stimulation as a positive control for a robust inflammatory trigger (Figure S5, Figure S8 b). By RNA sequencing, we found 211 RNAs unique for ORF8 besides 632 RNAs that reflect the inflammatory overlaps (Figure S 11 a) between the positive control LPS and ORF8 (Figure S11 b).

**Figure 3:**
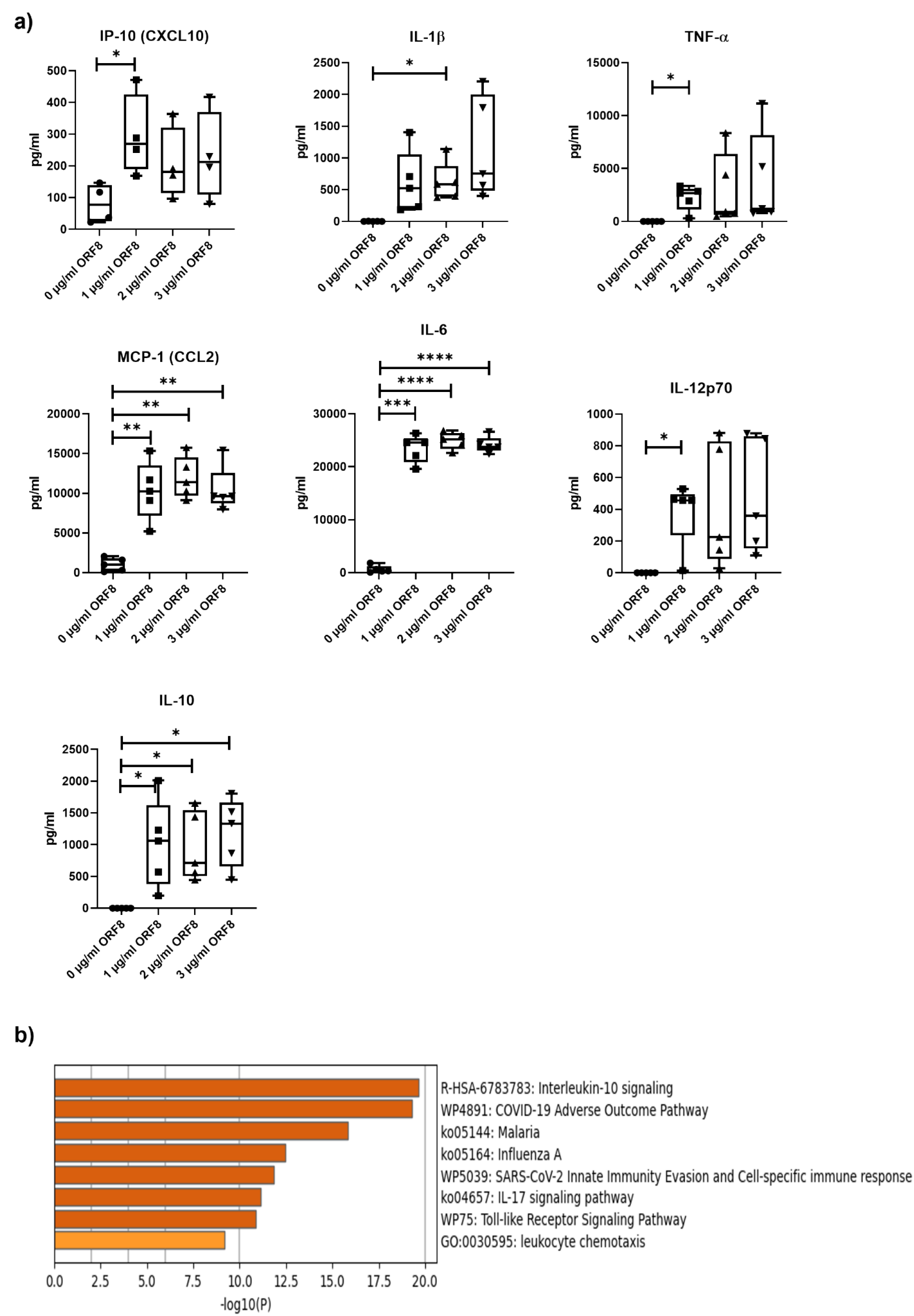
ORF8 induces a strong cytokine storm during dendritic cell differentiation. a) Monocytes were differentiated to immature dendritic cells by IL-4+GM-CSF in the absence (0 ng /ml) or presence of ORF8 in increasing concentrations (1 – 3 µg/ml). After 5 days, the supernatant was collected, and cytokine and chemokine levels were determined using a cytokine bead array (n = 5). IP-10: interferon-gamma induced protein 10 kD, CXCL10; IL-1β, interleukin 1β; TNFα, tumor necrosis factor α; MCP-1, monocyte chemotactic protein 1; IFNγ, interferon γ; TGF-β1, transforming growth factor β1; IL, interleukin; (* = .05, ** = .001, *** = .0001), b) Enrichment bar graph of the pathways in which the upregulated proteins after ORF8 treatment are involved; −log10(P): −log10 p-value

Using the metascape platform for enrichment and pathway analysis (28), we could show that this cytokine profile is significantly associated with COVID-19 adverse outcome pathways as well as SARS-CoV-2 innate immunity evasion and cell-specific immune response pathways (Figure 3 b).

To get deeper insights into the function and underlying mechanism of ORF8, we performed RNA sequencing of DCs that were differentiated in the presence or absence of ORF8 (120 h). We found a total of 843 regulated genes (p-value <0.05 after Benjamimi Hochberg correction) when comparing immature DCs with DCs, which received ORF8 during the differentiation (Figure 4 a, each group n = 4). Subsequent pathway and process enrichment analysis (metascape platform) of the ORF 8 activated DC RNA profile revealed a strong overlap with inflammatory and immune-activating pathways (Figure 4 b, S7 a). Next, we compared the regulated genes in ORF 8 treated DCs with regulated antiviral-related genes and cytokine genes in host cells after SARS-CoV2 infection (29). Here we found overlaps with the genes LAMP3, TRIM14, IL1A, CXCL8, IL15, IL6, CCL5, EBI3, LIF, and IL1B (Figure S7 b: host antiviral-related genes/host cytokine genes (SARS-CoV-2 infection)). In comparison with the expression profile of blood antigen-presenting cells in severe COVID-19 (30), we found an overlap with the genes CXCR4, TNAP3, KLF4, LICH, NR4A1, RETN, and IL1B (Figure S7 b: genes in APCs (SARS-CoV-2 infection)). All detected genes are related to severe infection. Using the Coronascape network (www.metascape.org/COVID) (28), we compared our data with single-cell mRNA sequencing data from monocytes (31) and bronchoalveolar immune cells (32) in patients with COVID-19. These cell types are the closest related cell populations available in the metascape databases. We found a great overlap in the gene profiles between our study and the published studies (Figure 4 c, d) and even a stronger overlap in genes that reflect the same pathways and function (Figure S7 a). Comparison of the functional enrichment analysis shows an overlap of the studied genes of all three studies mainly in the clusters of regulation of cytokine production, myloid leukocyte activation and leukocyte migration (Figure 4 c, Figure S7 a).

**Figure 4:**
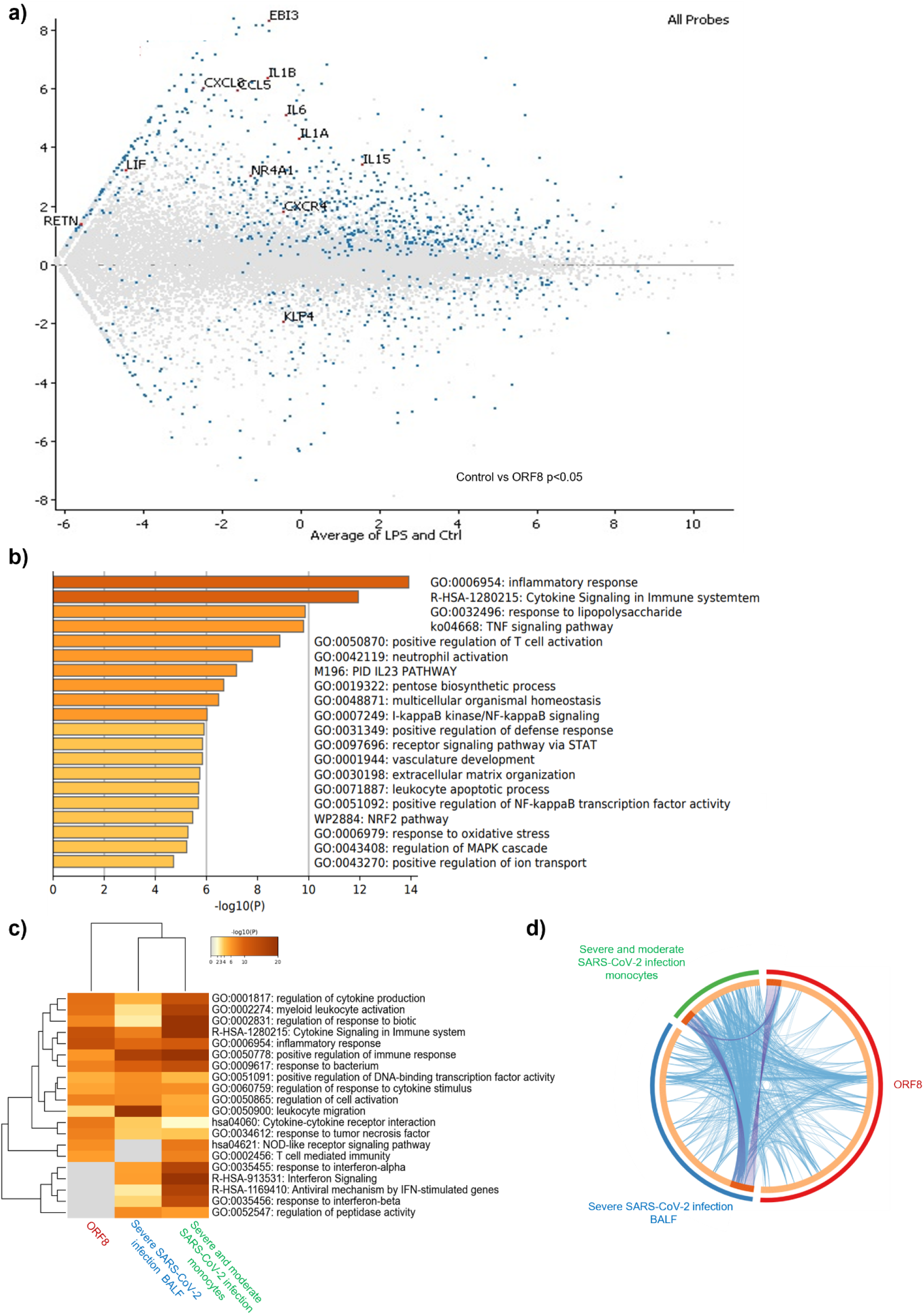
ORF8 induces an inflammatory mRNA profile involved in SARS-CoV-2 infection. a) MA plot visualizing gene expression differences between control vs ORF8 (gray dots: not significantly regulated genes; blue dots: significantly regulated genes (p-value <0.05 after Benjamimi Hochberg correction)); b) Enrichment bar graph of the ORF8 single gene list pathway and process enrichment analysis; c) Cluster analysis of enriched terms across input gene lists, colored by p-values (https://metascape.org/)(28). ORF8 = dataset of this publication (120 h); Severe SARS-CoV-2 infection BALF(32); severe and moderate SARS-CoV-2 infection monocytes (31). d) Overlap between gene lists: purple curves link identical genes; blue curves link genes that belong to the same enriched ontology term. The inner-circle represents gene lists, where hits are arranged along the arc. Genes that hit multiple lists are colored in dark orange, and genes unique to a list are shown in light orange: calculation source www.metascape.org/COVID(28).

### Detection of ORF8 in lung and serum of COVID-19 patients

Since monocytes circulate in the blood stream and emigrate into the e.g. lung tissue upon inflammation to differentiate into DCs (6, 11) we analysed whether ORF8 is present in these compartments of COVID-19 patients. In hospitalized patients ORF8 could be detected in the plasma (Figure S4 a) at different time points of infection and high concentrations in sever cases of the infected lung tissue (Figure S4 b). Lungs of Covid19-positive patients showed varying acute and chronic changes in the tissue from hyaline membranes, activated pneumocytes and increased amounts of alveolar and interstitial histiocytes. Sometimes focal intraalveolar haemorrhage and some giant cells could be found. On the other hand fibrosis of a developing organizing pneumonia could be found. Locotypical cells like vascular endothelial and smooth muscle cells and respiratory epithelium of bronchi also stained against ORF8 (Figure S4 b, patient 1 and 2 (sever), patient 3 (milder)).

### Lack of ORF8 neutralizing antibodies in COVID-19 patients

As demonstrated, ORF8 showed a strong immunomodulatory function in our study, and we were interested if patients infected with the SARS-CoV-2 virus produce antibodies against ORF8. Therefore we performed a custom-made ELISA to detect ORF8 antibodies in the convalescent plasma of 64 COVID-19 patients early tested positive for SARS-CoV-2 by RT-PCR (0-90 days). We found eight patients highly positive for anti-ORF8 IgG antibodies (Figure 5 a). To confirm this finding, another set of plasma (n=104) was analysed for the content of anti-ORF-8 antibodies. Again, eight patients were highly positive for anti-ORF8 IgG antibodies (Figure 5 c). Here, seven out of the eight samples also had a very strong reaction toward the SARS-CoV-2 spike protein (Figure S12). Next, we compared the serum samples of the ORF8 high and low/negative patients regarding the expression of pro-inflammatory chemokines and cytokines. In line with our in vitro findings, we found a significantly increased expression of IP-10 (Figure 5B) (p-value < 0.05 after). An overview of all performed antibody, PCR and the neutralization assays on patient serum can be found in the supplementary table (Table S1) and in the Spike (RBD) ELISA (Fig S12 a).

**Figure 5:**
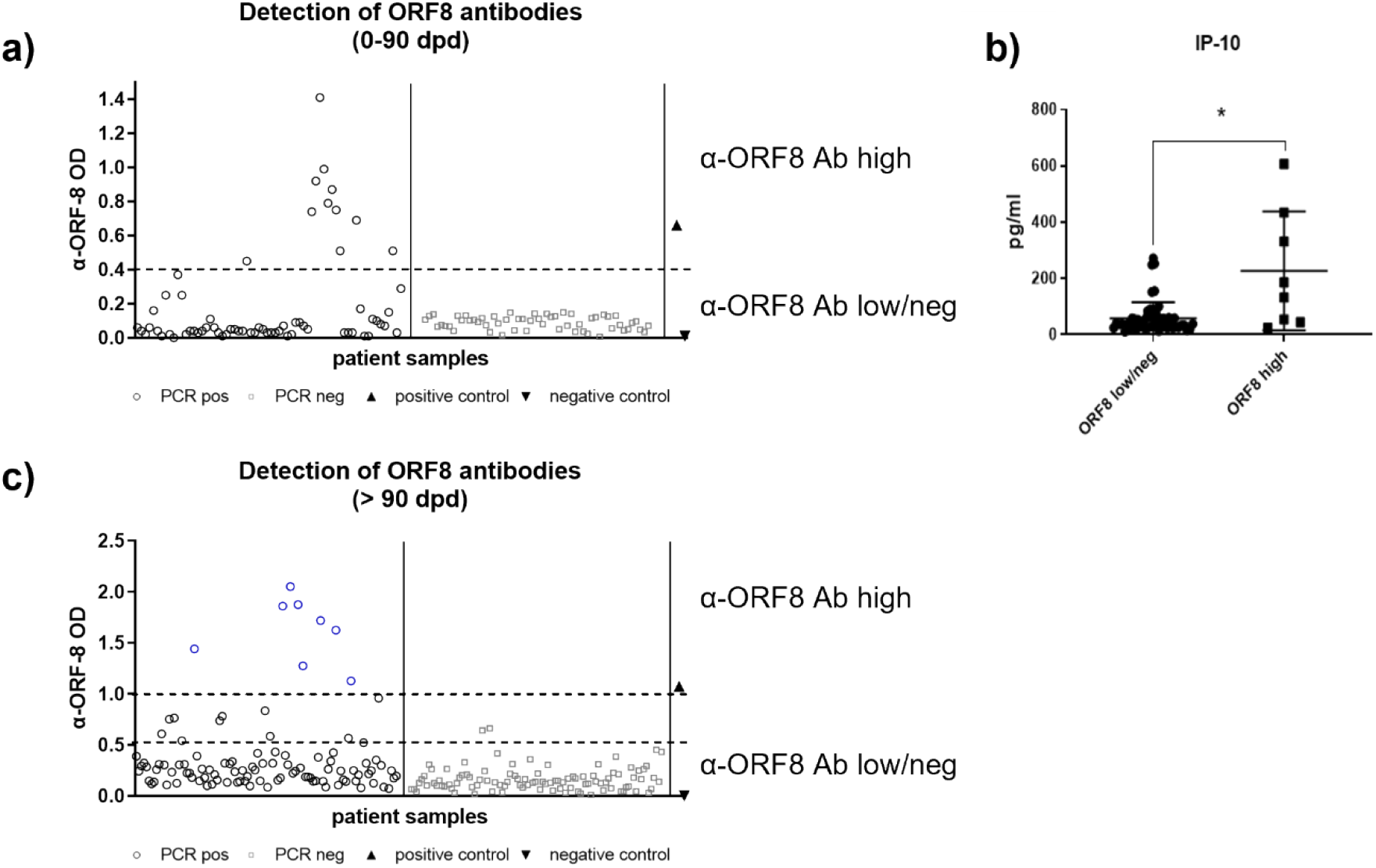
Detection of anti-ORF8 antibody titer in COVID-19 patients. a) Anti-ORF8 antibody levels of PCR positive for SARS-CoV-2 patients 0-90 days post-infection diagnosis (n=64; control: PCR negative patients; n=55). Anti-ORF8 antibodies levels were detected by an ORF8-ELISA; positive control: ORF8 positive plasma of a PCR positive COVID-19 patient; negative control: plasma of a healthy volunteer (PCR negative tested). ORF8 high: high titer of anti-ORF8 antibodies (ODs above 0.4); ORF8 low/neg: low titer of anti-ORF8 antibodies or negative for ORF8 antibodies (OD below 0.4). b) Samples from the same sera used in a) were used to compare the concentration of IP-10 between ORF8 high and ORF8 low/neg groups measured by a cytokine bead assay.c) Anti-ORF8 antibody levels of PCR positive for SARS-CoV-2 patients > 90 days post-infection diagnosis (n=104; control: PCR negative patients; n=100). ORF8 high: high titer of anti-ORF8 antibodies (ODs above 1.0); ORF8 low/neg: low titer of anti-ORF8 antibodies or negative for ORF8 antibodies (OD below 1.0). dpd: days post-diagnosis; OD: optical density; PCR pos: positive for SARS-CoV-2 detected by reverse transcription-quantitative PCR;

As we demonstrate, ORF8 shows a robust immunomodulatory capacity, and patients produce antibodies against ORF8; we analyzed if these antibodies from SARS-CoV-2 infected patients have the potential to neutralize the ORF8 effect on dendritic cells. Therefore, eight sera with high ORF8 antibody titer from the patient collective of late time points (Figure 5 c, labeled in blue) were analyzed. We pre-incubated the ORF8 protein with 5 µl of the serum samples (with correspond to an estimated anti-ORF8 antibody concentration between 1 – 5 µg) of each donor overnight to ensure the binding of the anti-ORF8 antibodies. The monocytes were incubated with the ORF8/sera complex or only serum during the differentiation. As a positive control, ORF8 without sera was used. We analyzed the ability of the sera to inhibit the binding and cyto-/chemokine production induced by ORF8. We were able to show that the antibodies directed against ORF8 could not reduce the binding of ORF8 to DCs (Figure 6 a). Only one of the donors (green box) had the capacity to neutralize this effect (Figure 6 a) and showed no enhanced binding of ORF8 to DCs (Serum 396, Figure 6 a). We analyzed this effect in more detail and seven out of the eight sera could not reduce the entire range in cyto-/chemokine production that is triggered by ORF8 (Figure 6 b). Analogous to our previous observation (Figure 2 b), we analyzed the surface expression of dendritic cell-specific markers and activation markers. We could show that the anti-ORF antibody-positive patient sera were not able to reduce ORF8 induced DC activation (Figure 6 c). Instead, we observed for most of the sera an enhancement of the ORF8 induced upregulation of MHC II, CD80, and CD83. By preincubating ORF8-Atto488 with anti-ORF8 IgG^+^ sera, we could show that ORF8 bound even better in combination with this sera. This effect could be reversed with blockade of the Fc receptor on the DCs (Figure S9).

**Figure 6.**
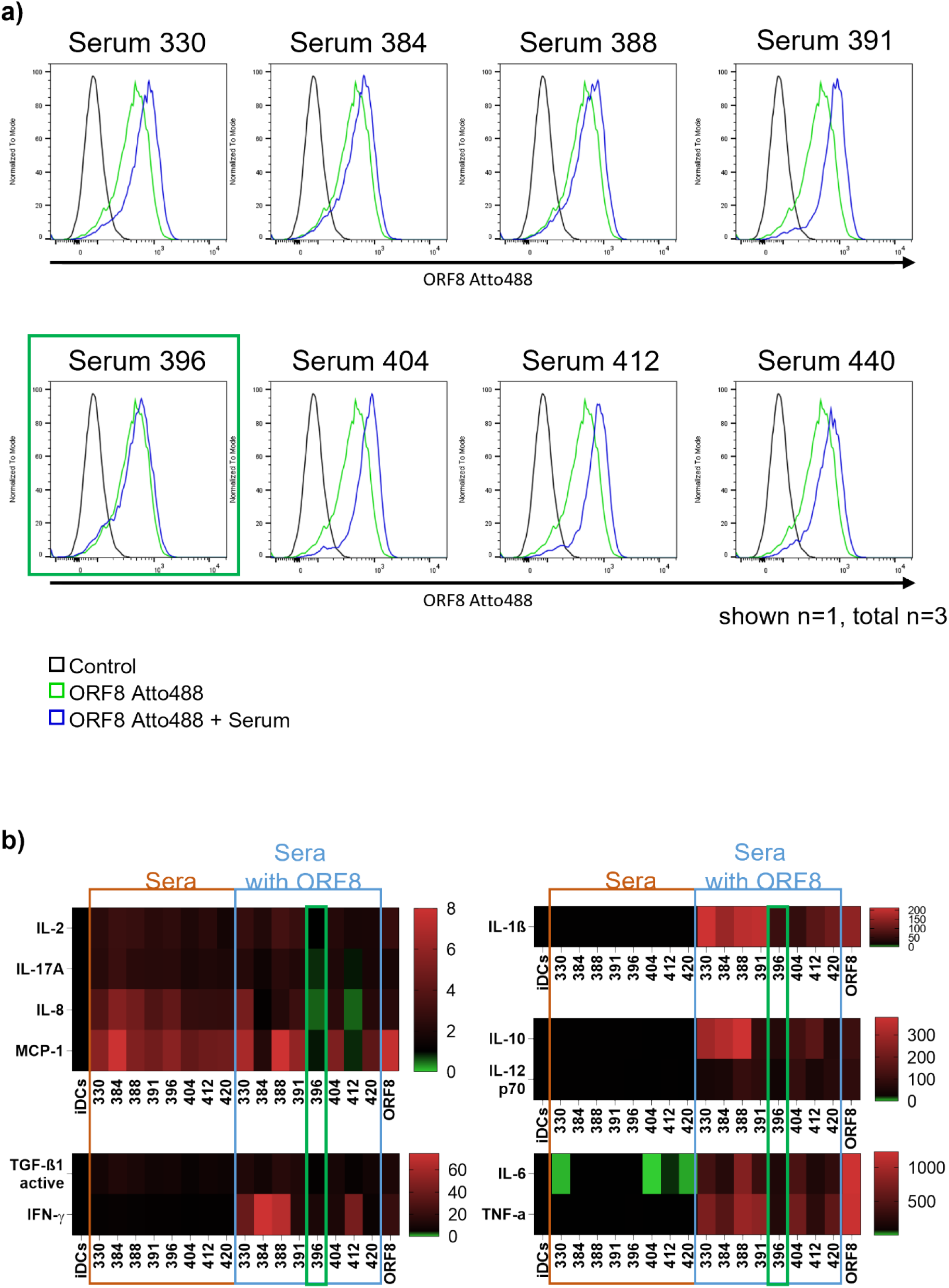
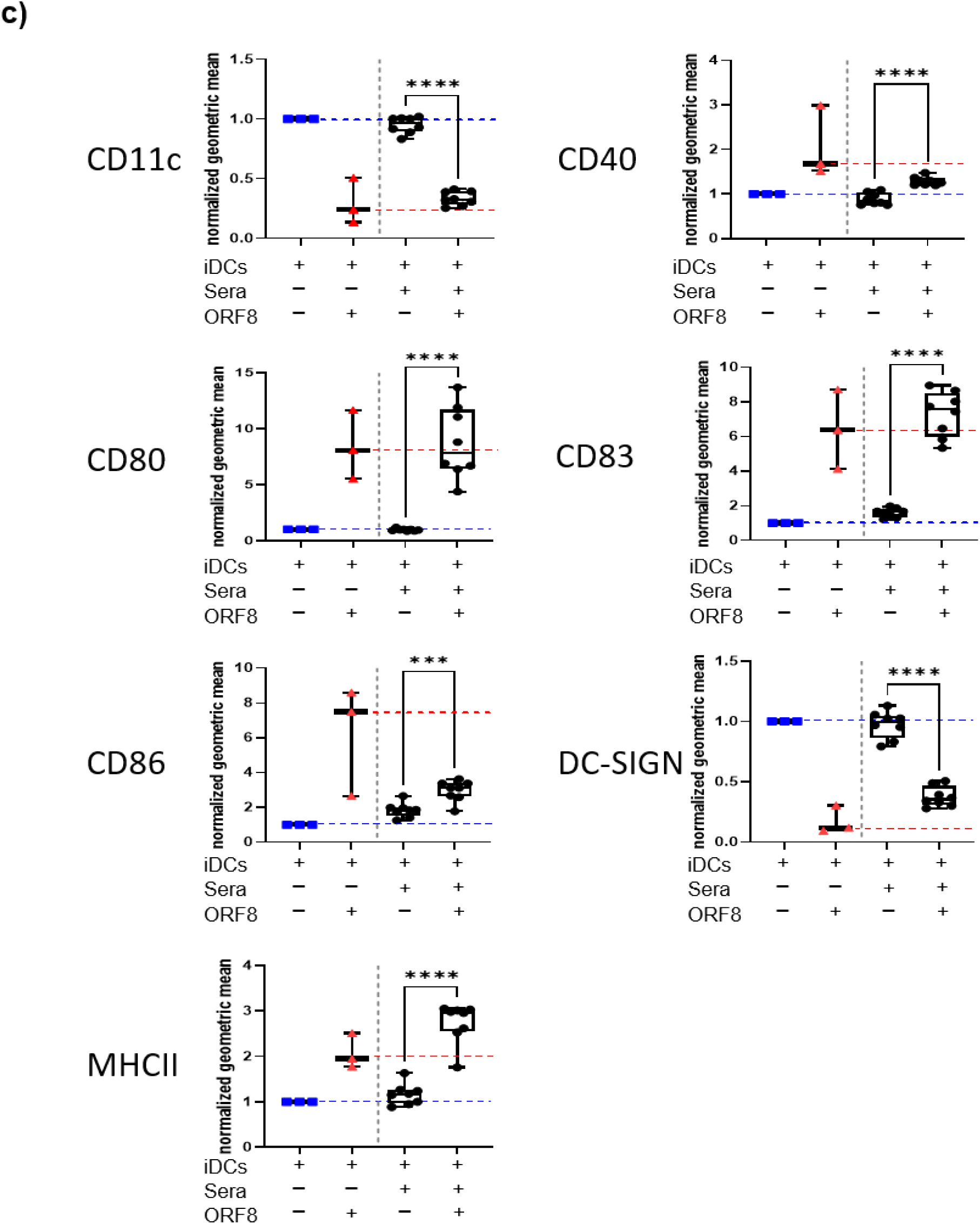
Failure of ORF8 serum antibodies to neutralize ORF8. a) The binding capacity of ORF8-Atto488 or ORF8-Atto488 in the presence of sera from anti-ORF8 positive donors was determined on immature DCs by FACS analysis. b) Multiplex Cytokine analysis of the supernatant of iDCs treated with either with ORF8 (n=3), patient serum alone (5 µl serum) (n=5) or a pre-incubated mix of ORF8 and 5 µl of serum (n=5). The sear was untreated to exclude any loss of antibody activities. Values of immature DCs were set to one, and expression changes were calculated accordingly and plotted in a heat map. c) Effect of anti-ORF8 antibody-containing sera on ORF8 mediated maturation of immature DCs. Immature DCs (iDCs) were incubated either with ORF8 (n=3), patient serum alone (serum) (n=5) or a pre-incubated mix of ORF8 and serum (n=5). Samples were normalized to the geometrical mean of immature dendritic cells.

In addition, in a small case series with sera from three patients hospitalized due to the severity of their SARS-CoV-2 infection, we found that, with the progression of the infection, the ORF8 induced cytokine storm even aggravated in the presence of anti-ORF8 IgG^+^ sera over time (Figure S10).

### Plasticity of the antibody recognized loop region of ORF8

The advantage of our study is that we used for the first time ORF8 expressed and purified from a mammalian system, whereas previous studies used bacterial-derived proteins (33). To compare the proteins and possible structural differences and a different potential structure for binding antibodies, we crystallized the eukaryotic expressed ORF8 protein and compared the structure against the bacterial expressed SARS-CoV-2 ORF8 protein and bat-related ORF8 variants.

The crystal structure of a SARS-CoV-2 ORF8 Asn78Gln mutant (ORF8_N78Q_) purified from HEK293 cells was determined to 2.6Å. The overall structure of ORF8_N78Q_ is comparable to the previously reported ORF8 (33) structure produced in *E. coli*, with some minor differences. The asymmetric unit is comprised of a single ORF8 monomer, with a second monomer related by a crystallographic 2-fold axis generating the biological dimer, linked by a disulfide bond at the interface (Figure 7 a). Each ORF8 monomer adopts an Ig-like fold comprising a pair of anti-parallel β-sheets held together by three disulfide bonds. Interestingly, while the ORF8 dimer interface is essentially identical to the previously reported structure, our ORF8_N78Q_ structure shows a difference in the relative orientation of the two monomers within the covalent dimer, with a rotation of ∼20° seen between monomers (Figure 7 b), suggesting that the ORF8 dimer interface is somewhat dynamic, despite the extensive, disulfide-bonded interface. This rotation gives the ORF8_N78Q_ structure a more “closed” shape, bringing the surface loops and surrounding structure between β2-β3 and β7-β8 of either monomer closer, towards the dimer interface (Figure 7 b). In addition, several solvent-exposed loop structures between strands β2-β3, β3-β4, β4-β5, and β7-β8 (numbered according to Flower *et al.* 2021) adopt slightly different conformations compared to deposited structures (Figure 7 c). In particular, residues 63-77, which includes the SARS-CoV-2-specific 73YIDI76 motif postulated to mediate ORF8 oligomerization (33), were unable to be modelled due to lack of electron density in the region (Figure 7 a). This is consistent with a previous report describing significant structural plasticity in this loop structure (33).

**Figure 7.**
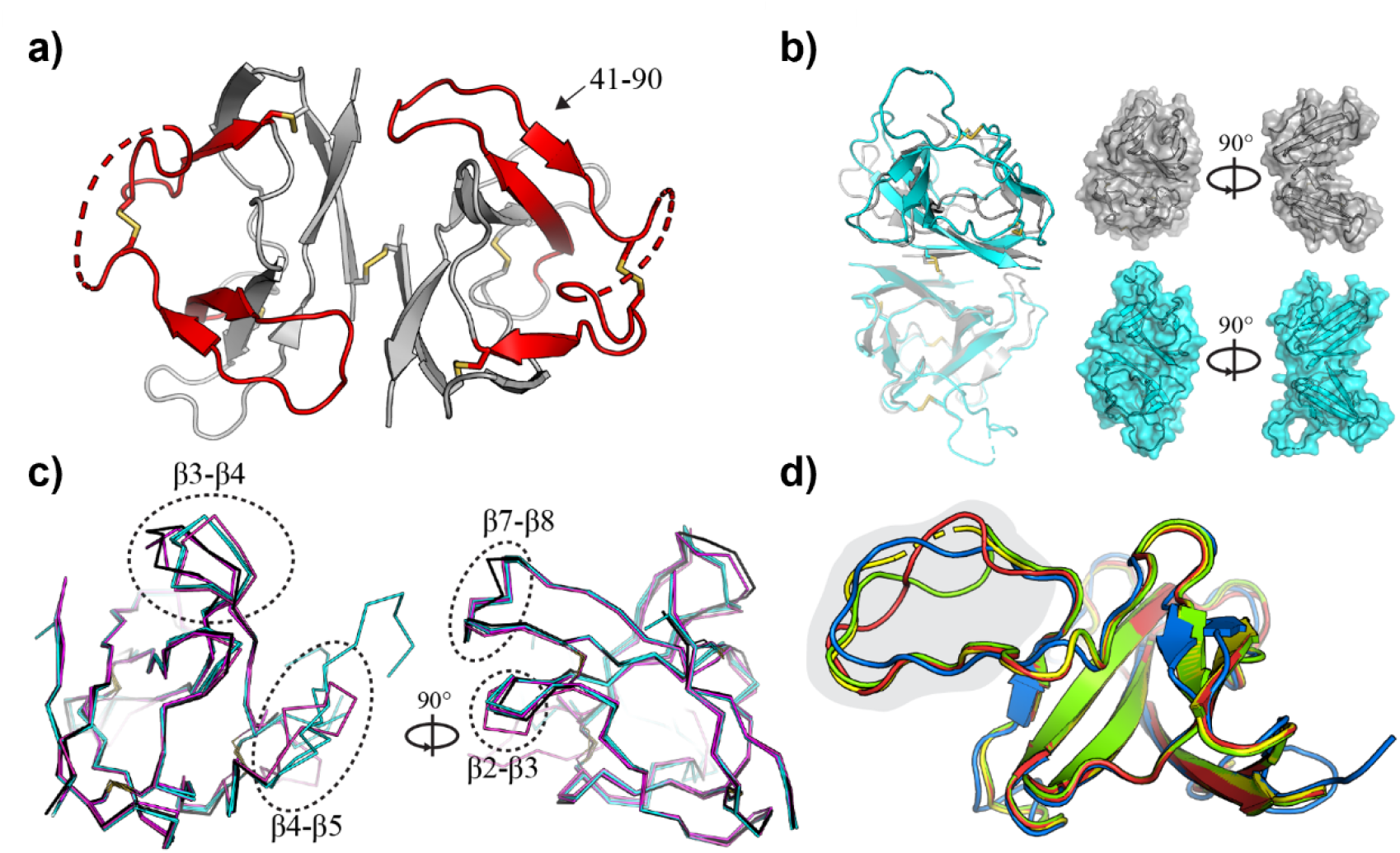
Crystal structure of SARS-CoV-2 ORF8 and related CoV homologues. a) Crystal structure of the SARS-CoV-2 ORF8 dimer. The sequence recognized by α-ORF8 antibodies (residues 41-90) (83) is highlighted in red and indicated. Intra- and inter-molecular disulfide bridges are shown as sticks, and regions with missing electron density (residues 63-77) are shown as dashed lines. b) Superposition of the ORF8 structure determined here (gray), and the previously deposited structure 7JTL (cyan). Chain A (transparent) of 7JTL and 7MX9 were aligned to show the change in the relative orientation of chain B within the dimer. Surface representations highlight the closed shape of 7MX9 relative to 7JTL. c) Individual monomers from 7MX9 (black), 7JTL (cyan), and 7JX6 (magenta) are superposed, and loop regions between β2-β3, β3-β4, β4-β5, and β7-β8 are indicated with dotted circles. Only Cα atoms are shown for clarity; d) Superposition of CoV ORF8 homology models. The bat-CoV RaTG13 (red), bat-like CoV Rs3367 (green) and SARS-CoV-2 (blue) ORF8 homologues, superposed on the SARS-CoV-2 ORF8 template (7JTL; yellow). Part of the ORF8-specific loop region (residues 63-77) is highlighted in gray. Disordered regions at the N-termini have been removed, and a single ORF8 monomer has been shown for clarity.

To further understand the structure of this loop region and possible consequences for the accessibility of antibodies, homology models were built for the ORF8 homologues of related CoVs (Figure 7 d). A superposition of the structural models of homologous ORFs from the bat-like CoV Rs3367, SARS-CoV and bat CoV RaTG13 with the SARS-CoV-2 ORF8 template highlights the potential structural plasticity of the region between different CoV species (Figure 7 d).

## Discussion

Recently, it was shown that ORF8 is secreted by SARS-CoV-2 infected cells and could be detected in the serum of COVID-19 patients (22). Furthermore, it was recently reported that a mutant SARS-CoV-2 strain found in Singapore displayed a deletion of 382 nucleotides in the region of ORF8 associated with decreased pathogenicity (34) (24) among several point mutations and smaller deletions (35–37), indicating the importance of ORF8 in the pathogenicity of this infectious disease. All main strands of SARS-CoV-2 contain the ORF8 protein (https://covariants.org/).

To study the patho-mechanistic role of ORF8 in COVID-19, we first investigated whether ORF8 is strongly expressed in infected tissue and is detectable in sera (Figure S4 b, a) as shown in the literature by Xiaosheng et al (38) and if ORF8 can directly bind to blood cells. PBMCs are composed of lymphocytes and monocytes, precursor cells for macrophages, and dendritic cells. In this study, we were able to demonstrate that extracellular ORF8 protein binds specifically to DCs (immature and mature, Figure 1 c, d, Figure S 3 b-d) as well as to its parental cells, monocytes (Figure 1 b, S3 a). Blocking ORF8 with a polyclonal α-ORF8 antibody significantly reduced its binding capacity (Figure 1 d), the isotype control antibody (Figure S3 d) was not able to reduce the binding and labled BSA (Figure S3 a, b) showed only a very low binding, demonstrating that the binding is specific and not a random protein interaction (Figure 1 d).

As ORF8 directly binds to monocytes, we were interested if ORF8 can also directly trigger the differentiation of monocyte to DCs. For several viruses, e. g. HIV-1 (39), it is known that their proteins are able to modulate dendritic cells and their function. Our findings show that ORF8 alone is not able to trigger differentiation since the precursor cells remained in a CD14^+^ monocyte stage (Figure 1 k). Since ORF8 independently was not able to modulate monocytes, we analyzed if ORF8 has an impact on the differentiation from monocytes to DCs. In fact, we could show that precursor monocytes differentiated by IL-4 and GM-CSF in the presence of the full-length ORF8 protein induced a pre-maturity in the differentiated DCs characterized by a mature cell morphology (40) as well as upregulated maturation makers MHCII, CD80, and CD40 as well as CD83 and CD86 (Figure 2 b). A dose dependent induction of the maturation can be observed (Figure S4 c, CD40 and CD 80 are shown). For further experiments we used the concentration that was in the range close to saturation and reflects most likely the strong expression in lung tissue as well as it is in line with the concentration used by other researchers like Lin X *et al.* (41) to study ORF8. We could exclude that protein contamination (Figure S1 b) or contamination with endotoxins is responsible for any effect on the cells. The amount of endotoxins in our protein preparation (Figure S1 c) were under the postulated levels of Harald Schwarz et al. that are needed to drive signaling and cytokine production in monocytes and DCs (42) and ORF8 showed an activation profile that differs from LPS stimulation (Figure S5). None of the other control proteins we used induced the activation and maturation of immature dendritic cells during differentiation (Figure S3 e).

Most interestingly, the dendritic cell marker DC-SIGN was downregulated entirely in the presence of ORF8 during differentiation (Figure 2 c). DC-SIGN was shown to be a receptor for SARS-CoV-2 on lung and kidney epithelial and endothelial cells (43). Even more interesting, it was also shown to be key for SARS-CoV-2 infection of DCs (44). From HIV infection, DC-SIGN is known to form an infectious synapse between infected DC and T cells that facilitates HIV infection (45). Furthermore, it was shown that DC-SIGN mediates the internalization of intact HIV into a low pH nonlysosomal compartment in dendritic cells. This internalization is needed for the trans-activation of T cells by DCs (27). In this study, we could show that blockade of ORF8 partly rescues the expression of DC-SIGN on the surface of immature DCs (Figure 2 d), pointing towards the interaction of ORF8 and DC-SIGN as described for HIV and DC-SIGN. In fact, we could further show that ORF8 interacts directly with DC-SIGN in a co-immune precipitation assay and that this interaction is reduced by an anti-ORF8 neutralizing antibody (Figure 2 e). Hence, DC-SIGN is a potential receptor for ORF8 and might be internalized upon binding to ORF8. DC-SIGN is known for its immune regulative role in DCs (46) and macrophages (47). A downregulation of DC-SIGN on the surface of DCs might lead to altered cytokine expression. It was shown that DC-SIGN ligation on dendritic cells results in ERK and PI3K activation and modulates IL-10 secretion (48).

ARDS is the main cause of death in patients infected with MERS-CoV, SARS-CoV, and SARS-CoV-2 (49, 50). ARDS is the final outcome of a cytokine storm reflected by key pro-inflammatory cytokines like IL-6, IL-8, IL-1β, GM-CSF, and chemokines such as CCL2, CCL-5, IP-10 (CXCL10), and CCL3 (51–53). Furthermore, high expression levels of IFN-γ, IP-10, IL-1β, MCP-1, and TNF-α have been detected in patients with SARS-CoV-2 infection (54). These inflammatory cytokines may activate the T helper function (Th1). Th1 activation is a key event in the activation of adaptive immunity. However, unlike SARS patients, patients with SARS-CoV-2 also have elevated levels of Th2 secreted cytokines such as IL-4 and IL-10, which belong to the anti-inflammatory panel of the body (50, 55, 56). Since DCs are a main source of cytokines, we analyzed the cytokine and chemokine fingerprints of cells that were differentiated in the presence of ORF8. We found a strong and significant upregulation of pro-inflammatory cytokines and chemokines IP-10, IL-1β, IL-6, IL-12p70, TNF-α, MCP-1, and IL-10 and type 2 cytokine (Figure 3 a). IFN-γ and IL-8 were upregulated by trend. The observed ORF8 profile showed a unique fingerprint and showed high similarities with COVID-19 adverse outcome pathways as well as SARS-CoV-2 innate immunity evasion and cell-specific immune response pathways (Figure 3B). Furthermore, the profile differed from LPS as a well-characterized pro-inflammatory control (Figure S5, Figure S11 b). The binding of ORF8 to monocytes was recently reported by other groups in preprinted articles (38, 57). Their observations back up and support our findings and demonstrate the unique function of ORF8 in the context of inflammation.

For further characterization of the pro-inflammatory response of the ORF8 treated DCs, we performed RNA sequencing. The pro-inflammatory and immune regulatory function of ORF8 could be confirmed by pathway and process enrichment analysis (Figure 4 b, c), and disease enrichment pointed out pneumonitis (Figure S11 a) as the highest reflection of our sequencing data. This result reflects very well the known phenotype of a SARS-CoV-2 infection. Our RNA dataset shows a great overlap (Figure 4 c, d) in inflammatory chemo-and cytokines of other studies (29, 30) and genes of the same functional group and pathways (31, 32) are regulated (Figure 4 c, d, Figure S 6 a, b) in the same direction in SARS-CoV-2 patient samples. From these data, we can conclude that ORF8 seems to play a role in the progression and course of the COVID-19 typical cytokine storm by activating DCs.

In addition to our in vitro findings, we could show in addition that patients positive for SARS-CoV-2 by PCR and high levels of anti-ORF8 antibodies have a significantly increased level of IP-10 early after infection (Figure 5 b). Recently, it was shown that plasma levels of IP-10 in COVID-19 patients are associated with disease severity and can be used as a predictive factor for the progression of the disease (58). In our two cohorts of convalescent plasma, about 10% of the SARS-CoV2 patients contained ORF8 IgG antibodies. Recently, it was shown that ORF8 antibodies could be detected in almost all SARS-CoV2 patients (59). Nonetheless, in accordance with our study, the cohort was divided into patients with high and low levels of ORF8 antibodies. Since the results of the study are waiting for validation by a conventional ELISA, further studies might explain the different findings.

Nevertheless, we could nearly not detect a neutralizing capacity of the identified anti-ORF8 IgG^+^ sera on the ORF8 induced cytokine storm. Instead, we found an enhanced binding of ORF8 pre-incubated with anti-ORF8 IgG^+^ sera to DCs (Figure 6 a). This enhancement seems to be mediated by the Fc-receptors on the dendritic cells since it could be blocked by conventional Fc-block (Figure S9). So, the production of anti-ORF8 antibodies seems to be contra-productive to inhibit the ORF8 mediated effect. Yajing Fu et al. discussed various mechanisms of SARS-CoV-mediated inflammation potential therapeutic tools to reduce SARS-CoV-2-induced inflammatory responses, including multiple methods to block FcR activation (60). In our study, we demonstrated that in most patients, antibodies against ORF8 cannot neutralize the immune-modulatory function of ORF8 (Figure 6 b, c) and even enhanced the binding (antibody-dependent enhancement (ADE)) of the protein to DCs (Figure 6 a, b, and c). For patients with severe SARS-CoV-2 infection in the intensive care unit, we even found that the effect gets enhanced with the progression of the disease (Figure S10). This effect is documented for several viruses like influenza (61), Dengue (62) as well as HIV-1 (63) (virus-induced ADE). ADE is intensively discussed for SARS-CoV-2 infection, and many studies demonstrated that antibodies against the virus were key to enhance the infection rate and are a critical challenge for developing further vaccines (64–66). In our study, we could show that the ORF8 protein induces in several patients a high antibody response (Figure 5) against the ORF8 protein and that these antibodies can bind to the FcR receptors (Figure S9) of dendritic cells and leads to an antibody-dependent enhancement (Figure 6 a). Therefore, an ORF8 neutralizing antibody or the use of FcR blocking antibodies could be beneficial for the outcome of severe SARS-CoV-2 infections. So, the detection of ORF8 antibodies in patient serum underscores the importance of our finding that ORF8 is an important factor in the development of COVID-19 and the missing neutralizing capacity of the serum antibodies shows that there is an unmet need to develop a neutralizing anti-ORF8 antibody to block, amongst others, the proinflammatory effect of ORF8 on dendritic cells. Since we used, for the first time, SARS-CoV-2 ORF8 purified from HEK293 cells that include all protein modifications like e.g. glycosylation, we determined the structure of this protein to a resolution of 2.6Å and observed that the overall structure of ORF8 is consistent with published structures of ORF8 isolated from *E. coli* (33). Interestingly, we noted a considerable degree of flexibility between monomers within the ORF8 dimer compared to reported structures, despite the presence of an identical dimer interface. To date, ORF8 has been implicated indirectly binding several cell surface receptors, including MHC-I (37), IL17RA (41), and DC-SIGN, described here. How the structural flexibility of ORF8 may be implicated in the binding and downregulation of multiple cell receptors to facilitate any immunomodulatory functions remains an open question. Similarly, ORF8 has been postulated to form large assemblies, mediated by SARS-CoV-2-specific sequence motifs (33). Whether ORF8 oligomerization is relevant with respect to DC-SIGN binding and DC cell differentiation remains to be determined.

We show that ORF8 causes the pre-maturation and activation of differentiating DCs and induces the secretion of pro-inflammatory cytokines. We further looked into the neutralizing antibody capacity of patient sera to neutralize the ORF8 induced effect. In conclusion, our findings show that ORF8 contributes to the cytokine and chemokine storm described for patients with SARS-CoV-2 infections (54, 67–69) by interaction with dendritic cells. Furthermore, our findings shed light on the function of ORF8 and its role in the development of ARDS.

Blocking of the cytokine and chemokine response mediated by the ORF8 protein might be an essential and novel additional step in the therapy of severe SARS-CoV-2 cases. Our findings showed that a polyclonal rabbit antibody against ORF8 is able to strongly reduce the binding of ORF8 (Figure 1 d) and the humoral immune response of dendritic cells (Figure S6). This demonstrates that the observed effects are a direct response to the ORF8 protein. Current treatment options focus on neutralizing single cytokines, chemokines (69), or the spike protein, but we could show for the first time evidence that neutralizing ORF8 and its strong pro-inflammatory characteristics is a promising approach to take out one of the possible key players that cause the cytokine storm. Kreer et al. (70) recently reported the longitudinal isolation of potent S protein neutralizing antibodies from COVID-19 patients. This strategy might also be promising in identifying ORF8 neutralizing antibodies. As antibodies failed to prevent or even enhance ORF8 binding to DCs (Figure 6), an experimental structure of ORF8 isolated under non-denaturing conditions confirms the stoichiometry of ORF8 and the biological relevance of the isolated disulfide-linked homodimer relevant structure is important for the further development of inhibitory antibodies targeting ORF8. Unfortunately, it is difficult to ascertain how the flexibility of ORF8 would influence antibody binding in the absence of an experimental structure of ORF8 bound to an antibody of interest. The apparent flexibility of this region suggests that it may adopt alternate conformations upon binding to an interaction partner. As suggested elsewhere (33), this region of ORF8 may also contribute directly to the formation of higher-order oligomers, although the relevance of such structures has not been demonstrated directly. In the event that such structures are functionally important for SARS-CoV-2 pathogenesis, antibody-mediated disruption of these complexes may be important.

In general, we conclude that ORF8 via binding to dendritic cells is an important player in SARS-CoV-2 induced chemokine and cytokine storm in patients with severe outcomes and that targeting ORF8 may open new treatment avenues for severe COVID-19 patients (graphical abstract Figure S12).

## Methods

### Protein design, expression, and purification

The ORF8 open reading frame was cloned into a sleeping beauty transposon vector either with or without a Strep-tag at the N-term (Figure S1 A). HEK293 cells were transfected with the ORF8 plasmid together with the transposase coding plasmid (1/10) with Fugene HD (Promega, Germany) and after one day selected with 3 µg/ml puromycin for three days. The cells were then transferred to Triple Flasks in DMEM/F12 supplemented with 10% FCS and the cells were grown to confluence. Protein expression was induced with 1 µg/ml doxycycline in DMEM/F12 medium and 2% FCS. Every second or third day, the medium was collected and replaced. The collected supernatants were purified with Streptactin XT sepharose (IBA Lifesciences, Germany) according to the manufacturer’s protocol. Next, the recombinant proteins were dialyzed against phosphate buffered saline (10 mM Na_2_HPO_4_ 2H_2_0, 1.8 mM KH_2_PO_4_, 140 mM NaCl, 2.7 mM, pH 7.4) or bicarbonate buffer (50 mM NaHCO_3_, 500 mM NaCl, pH 8.3). Protein purity was analyzed by SDS-gel and staining with brilliant blue (Figure S1 B).

### Labeling of ORF8 with Atto 488 dye

Where indicated, Atto 488 NHS ester dye (Attotec, Germany) was dissolved in DMSO and added in 3 times molar excess relative to the protein concentration. The labeled proteins were dialyzed 3 times against PBS buffer.

### Detection of endotoxin in the purified ORF8 protein

For the determination of endotoxin levels in the purified ORF8 protein solution, a Chromgenic Endotoxin Quant Kit (Pierce/Invitrogen, Germany) was used according to the manufactureŕs instructions. The detection range was from 0.1 EU/ml to 1 EU/ml for the standard curve. All measuring points were performed in independent triplicates and adjusted to background subtraction. The measurement (Epoch reader (BioTek, Germany)) and data analyzes were performed with the Gen5 software (BioTek, Germany). From the standard curve, the values of ORF8 and BSA endotoxin levels per µg protein were calculated and applied as data points to the standard curve.

### For all presented experiments and ethical approvals

The n number represents biological replicates with a minimum of n = 3, including a blind study of different donors with ethical review (N°11-339) and (DRKS00021468 EIKOS) of the ethical comity university hospital cologne. For the study consent from all participants was obtained. For the statistical analysis the software GraphPad Prism version 9 GraphPad Software, San Diego, USA) was used and a one-and tow-way ANOVA statistic was performed.

### Polyclonal antibody against ORF8

Recombinant ORF8 was used to immunize two rabbits. After two months, sera were collected and the antibodies affinity purified via an ORF8 affinity column. Tag free ORF8 was coupled to CNBr-activated Sepharose (GE Health-care), and after applying the sera to the column the specific antibodies were eluted with 150 mM NaCl, 0.1 M triethylamine, pH 11.5 and neutralized with 1M Tris-HCl, pH 6.8. The two antibodies were tested by Western blot analysis. 25 µl of the HEK293 supernatants were applied to a 12% SDS-PAGE gel under reducing conditions and visualized either by Coomassie Blue staining or by Western blot analysis utilizing the ORF8 affinity-purified antibodies from rabbit 1.

### Isolation and differentiation of monocytes to dendritic cells

Mononuclear cells (PBMCs) were freshly isolated using a density gradient (Lymphoprep, the density of 1.077 g/ml, Stemcell, France) from leukocyte concentrate (leukapheresis apheresis chamber). Ethical clearance for the use of human material was obtained from the ethical review committee of the university hospital Cologne (N°11-339). The harvested PBMCs were washed with PBS and resuspended in X-Vivo 15 (Biozym, Germany). The viability and cell number were obtained by trypan blue staining and counting. For the enrichment of monocytes, we used the adherence strategy as described before (71). For each well of a 6-well plate, we seeded 2 x 10^7^ PBMCs in 2 ml of X-Vivo media. The cells were incubated for 75 min at 5% CO_2_ and 90% humidity at 37°C to allow adherence of monocytes. The non-adherent fraction (NAF) was removed from the 6-well plates by intensive washing. For differentiation to immature moDCs, X-Vivo 15 media was supplemented with 100 ng/ml human granulocyte-macrophage colony-stimulating factor (GM-CSF) and 20 ng/ml human interleukin 4 (IL-4) (both cytokines obtained from Peprotech, Germany), and cells were cultured with 5% CO_2_, and 90% humidity at 37°C for five days. To analyze the function of ORF8 (used amount indicated in the figures), GM-CSF and IL-4 got replaced by ORF8 protein or ORF8 as well as Lipopolysaccharide (LPS; positive control) was used as an add-on together with the differentiation cytokines.

### FACS analysis of ORF8 binding and of the analysis of monocytes and DCs

For the flow cytometry procedures, specific commercial antibodies (see below) were diluted in FACS buffer (1% BSA in PBS). Prior to staining, we performed an Fc block (BD Biosciences, Heidelberg, Germany) to avoid unspecific false-positive events for all stainings. For the direct interaction of ORF8 to the cells, they were pre-incubation with the ORF8-Atto488 protein before the staining of all other cell surface markers. As a control for specificity and to exclude random protein binding we used BSA-Atto488 as a control staining. For the staining of monocytes, a combination of CD14, DC-SIGN, and CD1c (BioLegend, London, UK) was used. For the immature moDCs, we used MHCII, CD11c, CD40, CD80, CD83, CD86, and DC-SIGN (BioLegend, London, UK). The cells were stained for 30 min at 4°C. The cells were then washed twice in ice-cold FACS buffer, and acquisitions were performed on a FACSCanto II Flow Cytometer (BD Biosciences, Heidelberg, Germany). Warranty of the stability of acquisition was verified for every experiment using a dotplot Time *vs*. SSC-H or fluorescence. The data were analyzed with the FlowJo v10 software (BD, Vancouver, Canada).

### Blocking of ORF8 by polyclonal rabbit Ab

For blocking the labeled and unlabelled ORF8 protein, we pre-incubated the protein with an anti-SARS-CoV-2 ORF8 detecting antibody (Antibody.com, London, UK) for half an hour on ice to induce ORF8/antibody interaction. Due to the sodium azide in the antibody solution, we used the same µg of the antibody (max. 2 µg) as µg of the ORF8 protein. In addition, we used the antibody alone in the same concentration as in the complex as an additional control for cytotoxicity. As an isotype control an unspecific polyclonal rabbit antibody was used (NEB, Frankfurt, Germany). To block ORF8 during differentiation it was preincubated with the polyclonal rabbit anti-ORF8 antibody or isotype over night and added to the monocytes (t=0) at the same time as GM-CSF and IL-4 for the differentiaton.

### Co-immunoprecipitation of ORF8 and DC-SIGN

For the co-IPs 25 µl of streptavidin-coated magnetic beads (NEB, Frankfurt, Germany) were used per separation. The beads were washed with 500 µl PBS once before use and resuspended in 50 µl PBS. For immune precipitation (IP), beads were pre-incubated at 4°C for 1h with agitation with empty control (no protein) or 2 µg ORF8 protein with Strep II tag or 2 µg ORF8 plus 2 µg of an anti-ORF8 antibody (Antibody.com, London, UK). Unbound protein was washed from the beads with 500 µl PBS, spun down and resuspended in 50 µl PBS. For protein-protein interaction, pre-incubated beads were incubated a second time with 2 µg of a recombinant human DC-SIGN protein with a human Fc tag (Sino Biological, Köln, Germany) for 1 h at 4°C with agitation. The beads were washed three times with 500 µl PBS before denaturation and separation by a polyacrylamide gel (4-12% gradient). The gels were blotted to a nitrocellulose membrane and blocked with 5 % milk in TBST buffer. For the staining, a direct HRP conjugated anti-human Fc (Invitorgen, Wiesbaden, Germany) was used, and bands were detected with an imager (Bio-Rad, Feldenkirchen, Germany).

### Immuno blot and immuo histochemistry

Circulation of the ORF8 was analyzed in sera of five hospitalized patients collected at two timepoints (t1 and t2). Therefore, 2 µl of sera was boild in 22 µl dendaturing laemmli loading buffer and seperated with a 4 – 12% NuPage SDS gel, running with MES buffer. As a control healthy donor sera was spiked in with bacteria expressed ORF8 (RP-87666; Invitrogen, Darmstadt, Germany) with the indicated amount/ml. The seperated protein was transferred to a nitrocellulose membrane. The membrane was blocked with 5% milk for one hour, followed by the primary antibody 1:1000 (anti-ORF8 C-term (ABIN6992307), antibodies-online.com, Aachen, Germany) over night. As a secondary antibody we used an F(ab’)_2_ anti-rabbit HRP 1:7000 (NA9340, Amersham, Freiburg, Germany). The development was performed with West Pico PLUS Chemiluminescent Substrate (Thermo, Darmstadt, Germany) with an imaging system from bio-rad (Bio-rad, Feldkirchen, Germany).

For immunohistochemistry ORF8 was detected in formalin-fixed, paraffin-embedded sections (3 µm thick) of SARS-CoV-2 infected patients after citrate epitope retrieval (pH 8), using an anti-ORF8 monoclonal antibody (Monoclonal Mouse IgG_2A_ Clone # 1041422, R&D, Germany) in a BOND immunostainer at a dilution of 1: 500. After blocking, incubation with H2O2 for 5 min and enhancer for 10 min the antibody was developed using poly-HRP-anti mouse/rabbit IgG (Bright Vision, Medac-diagnostica, Germany) and DAB away kit (Biocare Medical, Germany). In addition, the morphology of the Formalin-Fixed Paraffin-Embedded (FFPE) tissues was examined using routine H&E staining.

### Multiplex analysis of cytokines

We collected supernatant from differentiated DC of several donors in the presence or absence of ORF8 protein. The supernatant was cleared twice by centrifugation to exclude cells and cellular fragments before storage at −80°C. For the LegendPlex assay, cell supernatant was thawed on ice. We followed the manufactures instructions of the LEGENDplex™ HU Essential Immune Response Panel (BioLegend, San Diego, USA). The acquisitions of the beads were performed on a FACSCanto II Flow Cytometer (BD Biosciences, Heidelberg, Germany). The data were analyzed with the LEGENDplex v8 software (BioLegend, San Diego, USA) to determine the standard curves and current concentrations of the samples. The data were transferred to GraphPad Prism v9 (GraphPad, San Diego, USA) for visualization and statistical analysis. For the statistics, a one-way ANOVA analysis was used, and significance was marked with one to four stars.

### Patients and sample collection

COVID-19 patients with respiratory samples positive for SARS-CoV-2 by reverse transcription-quantitative PCR (RT-qPCR) after admission to the University Hospital Cologne were included in this study. Our study enrolled a total of 268 adult patients, based on recruitment of available patients with a positive COVID-19 test between day zero and ninety days after the first diagnosis to cover the early onset of the infection. Ethical clearance for the use of human material was obtained from the ethical review committee of the University Hospital Cologne. For 10 patients, information about age and sex, as well as hospitalization, were not available. In the study cohort, the age ranged from 24 to 80 years, with an average of 41. The study included 37.5 % male, 46,875 % female and 15,625 undefined individuals. Twelve patients were hospitalized with a follow-up, and for 76 % of the patients neutralizing antibodies for the virus were detected (DKRS number: DRKS00021468 EIKOS).

For the observation of potential inhibitory ORF8 antibodies in patients that were hospitalized, we included three patients with ages 51, 58, and 75. The patients were hospitalized and treated with oxygen and symptomatic treatment and two serum time points with a range between two and five days were collected. For one patient, we had a longer serum follow-up period of 11 days with four time points. This patient got a High-Flow oxygen, Remdesivir, and Dexamethason treatment.

### ORF8 ELISA

For the ORF8 IgG ELISA assays, 96 well Maxisorp Nunc Immuno plates (Thermo Fisher, Denmark) were coated over-night with 0.5 µg ORF8 in 50 µL TBS, pH 7.4 at 4° C. After washing with TBS, unspecific binding sites were blocked at room temperature (RT) with 3% BSA/TBS blocking buffer for 1 h. Plasma samples were diluted 1:100 with blocking buffer and duplicates of 50 µL were measured. After 90 minutes of incubation (RT) the ELISA plates were washed 3 times with TBS buffer and a rabbit anti-human HRP antibody 1:3000 (Dako) in blocking solution was applied for 1 hour (RT) and the plates were washed 3 times. Horseradish peroxidase activity was detected with 50 µL 1-Step Ultra TMB ELISA substrate solution (Thermo Fisher). Then the reaction was stopped with 50 µL 10% H_2_SO_4_ and the absorbance was measured at 450 nm. The second set of samples were measured by the Immundiagnostik AG using a newly established ELISA for the detection of ORF8 IgG based on the above described ELISA (Figure 5) (72). The immunoassay detects human anti-SARS-CoV-2 ORF8 IgG isotype antibodies. The manufacturer provides positive and negative controls and a cut-off sample to facilitate the evaluation of the patients. Due to the fact that in the negative group (n=100, PCR negative) some of the plasma from patients with an infection showed a low reactivity towards ORF8, the cut-off had to be set at OD= 1.1. The bindings were plotted and analyzed with GraphPad Prism 8 and 9.

### Structural determination of ORF8_N78Q_ and modeling of CoV ORF8 homologues

Purified ORF8_N78Q_ was loaded onto a Superose 6 (GE Healthcare) gel filtration column and eluted in 20 mM Tris pH 7.5, 150 mM NaCl. The eluted protein was concentrated to 20 mg/mL and screened in 96-well format via sitting-drop vapour diffusing using an NT8 (Formulatrix) robotic drop-setter. A single crystal was identified in well A4 of a PEG/Ion crystallization screen (Hampton Research) with well solution containing 0.2M LiCl, 20% (w/v) polyethylene glycol (PEG) 3350. A single crystal was swept through a drop-containing well solution supplemented with 20% glycerol and mounted directly into the cryostream. X-ray diffraction images were collected using a Rigaku 007-HF MicroFocus X-ray generator and an R-AXIS IV++ image plate detector. X-ray diffraction images were integrated and scaled using XDS (73), and merged using Aimless (74) within the CCP4i2 software package (75). Initial phase estimates for the ORF8_N78Q_ structure were obtained by molecular replacement within Phaser (76), using a modified polyalanine search model derived from PDB entry 7JTL (33). An initial model was build using AutoBuild (77), and successive rounds of model building and refinement were performed using Coot (78) and phenix.refine (79), respectively. Coordinates and structure factors for the ORF8_N78Q_ have been deposited to the Protein Data Bank under PDB ID 7MX9, and X-ray data reduction and refinement statistics are listed in Table 1. Structural models for additional CoV ORF8 homologues were built using Modeller (80) with sequences acquired from NCBI accession numbers AGZ48826.1 (bat SARS-like CoV Rs3367), AHX37565.1 (Bat-CoV RaTG13) and AAU04641.1 (SARS-CoV). All structures were modeled using chain A of the previously reported and deposited ORF8 structure 7JTL (33) as a template.

**Table 1:** Data collection and refinement statistics

|  |  |
| --- | --- |
| **Crystal** | ORF8<sub>N78Q</sub> |
| **X-ray source** | Rigaku MicroMax-007 HF |
| **Crystal geometry** |  |
| Space group | P4<sub>2</sub>2<sub>1</sub>2 |
| Unit cell (Å) | $a = 51.35$ $b = 51.35$ $c = 74.09$;<br>$\alpha = \beta = \gamma = 90^\circ$ |
| **Crystallographic data** |  |
| Wavelength (Å) | 1.541870 |
| Resolution range (Å) | 42.20 – 2.60 (2.72 – 2.60) |
| Total observations | 43786 (5496) |
| Unique reflections | 3379 (400) |
| Multiplicity | 13 (13.7) |
| Completeness (%) | 100 (100) |
| $R_{\text{merge}}$ | 0.166 (0.796) |
| CC1/2 | 0.997 (0.932) |
| $I/\sigma I$ | 11.8 (2.6) |
| Wilson B-factor (Å<sup>2</sup>) | 42.18 |
| **Refinement statistics** |  |
| Reflections in test set | 336 |
| Protein atoms | 1436 |
| Solvent molecules | 11 |
| $R_{\text{work}}/R_{\text{free}}$ | 0.2432/0.2966 |
| **RMSDs** |  |
| Bond lengths/angles (Å/°) | 0.004/0.748 |
| **Ramachandran plot** |  |
| Favored/allowed (%) | 94.05/5.95 |
| **Average B factor (Å<sup>2</sup>)** |  |
| Macromolecules | 43.08 |
| Solvent | 34.80 |

### RNA sequencing and analysis

RNA sequencing was performed in collaboration with the Cologne Center for Genomics. The Illumina® TruSeq® mRNA stranded sample preparation Kit was used for library preparation. Briefly, mRNA was purified and fragmented from 1µg total RNA after poly-A selection. After reverse transcription using random primers, second strand cDNA synthesis was performed and, after end repair and A-tailing, indexing adapters were ligated. Purified and amplified products were used to create the final cDNA libraries. Automated electrophoresis (Agilent 420 tape station) was applied for validation and quantification. Equimolar amounts of the library were pooled and quantified by the Peqlab KAPA Library Quantification Kit. Paired end sequencing with 100bp read length was performed with the Illumina novaSeq 6000 instrument. RNA sequencing reads were aligned using the HISAT2 alignment program^1^ (81), and data were processed and analyzed using SeqMonk V1.42.0 software (https://www.bioinformatics.babraham.ac.uk/projects/seqmonk/) and FunRich V3.1.4 (http://www.funrich.org/) (82). For further analysis and comparison with COVID-19 datasets, we used the metascape platform (Database 2021-05-01, https://metascape.org/) published by Zhou et al. (28).

## Acknowledgments

We thank Semra Oezcelik for her excellent technical assistance. We owe special thanks to Dr. Franz Paul Armbruster, Claudia Schumann, Sabine Friedl, Dagmar Szellas, and Dr. Eva-Maria Rogg from Immundiagnostik for establishing the ORF8 ELISA kit in collaboration with the University Hospital Cologne.

We thank Gabriele Braun from the University Hospital Cologne for her assistance and help in handling the cell culture experiments. Thank you to Mert Mestanoglu for helping with the editing.

## Funding

This work is supported by German Research Foundation (DFG) research unit FOR2240 (www.FOR2240.de) BO4489/1-1, BO4489/1-2, BO4489/3-1 (F.B.) Cu 47/9-1 and 12-1 and CMMC Cologne (FB, CC, M.K.); German Research Foundation (DFG) research unit - FOR 2722 (M.K.).

## Methods: data sharing statement

X-ray structure and modelling data are available online and been deposited to the Protein Data Bank under PDB ID 7MX9.

For Sequencing data and LegenPlex data, please contact

All co-authors concur with the submission of the manuscript to the Journal. The material submitted for publication has not been previously reported and is not under consideration for publication elsewhere. In addition, the authors do not have any conflicting financial interests.

## In Memoriam

***With this study, we would like to remember and honor all patients with severe SARS-CoV-2 outcomes. We hope that our study will help to ensure that their story is not forgotten and that it will be possible to prevent severe cases in the future. We thank all the helpers for their ongoing efforts and fight for each patient*.**

## Supplementary Information

**Figure S1:**
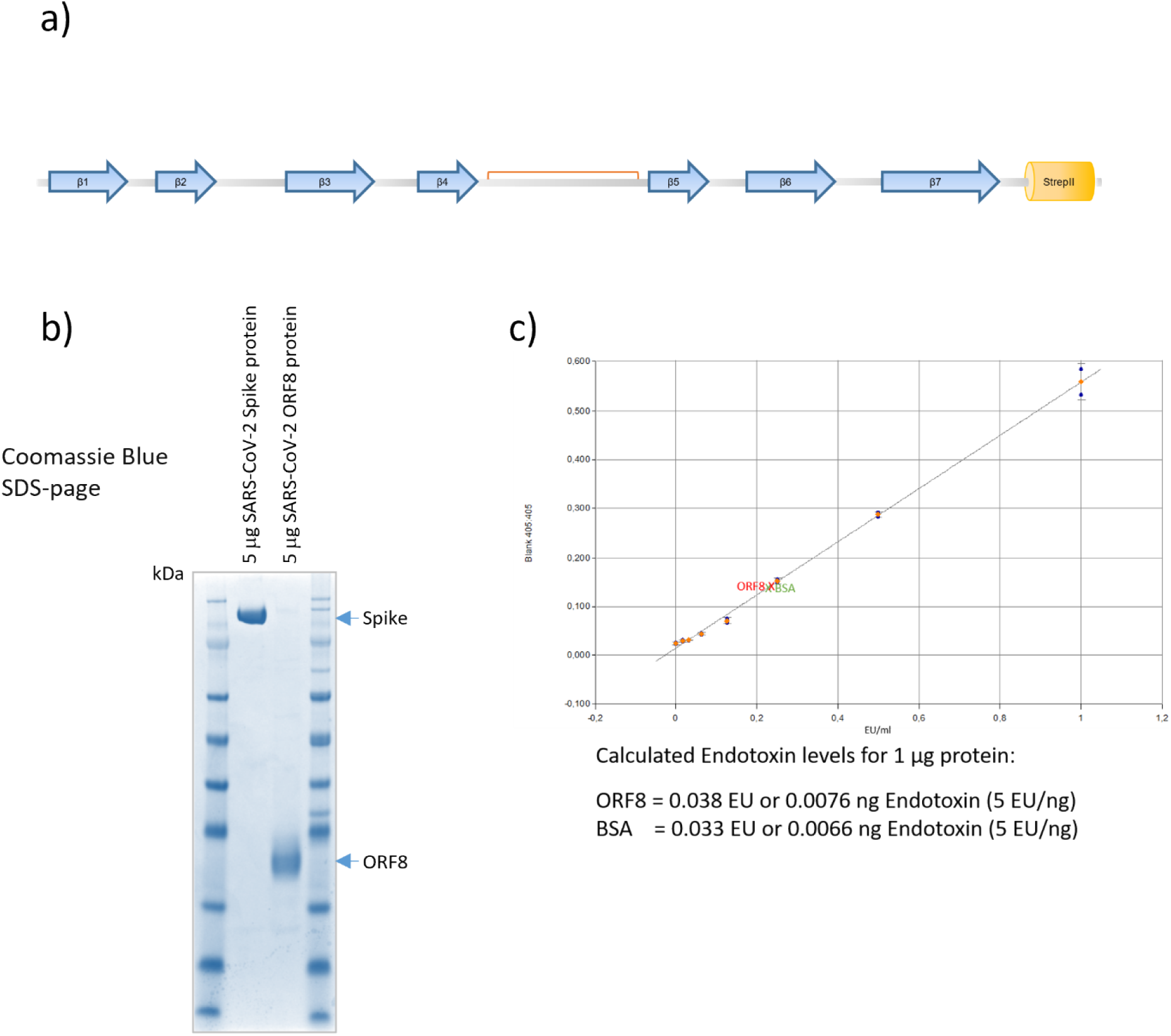
ORF8 protein structure, purification, and Endotoxin level. a) Secondary structure of the cloned SARS-CoV-2 ORF8 (beta-sheet structures are represented in blue arrows, adapted from Flower et al.). Putative disulfide bonds are marked in orange. The c-terminal tag (yellow) is the StrepII tag that was used for the purification. b) Protein purity analysis by SDS-gel and staining with brilliant blue. For each lane, 5 µg of protein were loaded. c) For the determination of Endotoxins, a LAL based Endotoxin assay was performed. The solid line represents the mean of the standard curve. In green and red, the detected values of the used BSA and ORF8 are shown.

**Figure S2:**
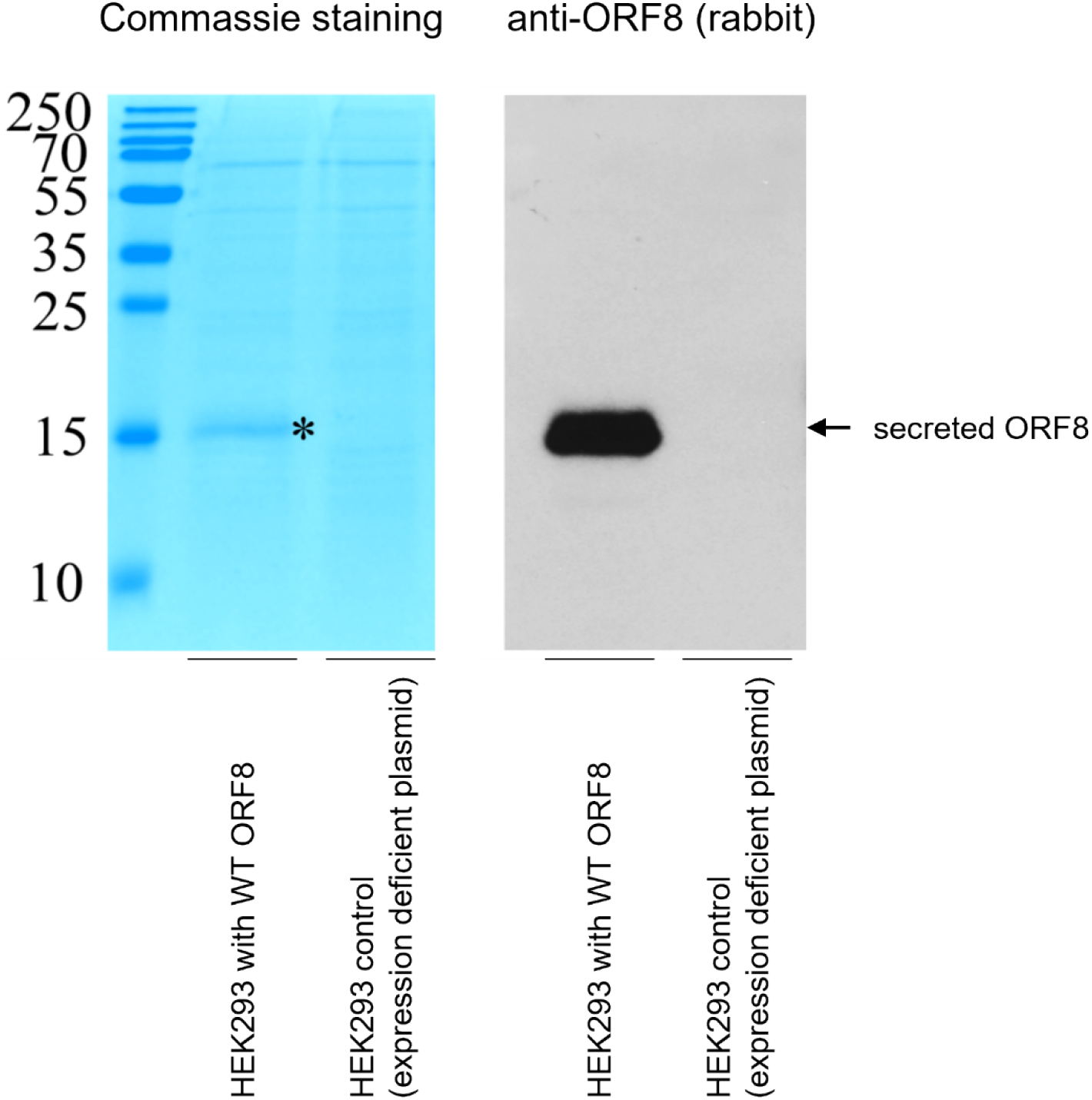
ORF8 protein is secreted into the supernatant in cell culture. a) Coomassie Blue staining of a SDS-gel loaded with supernatant of wild type ORF8 secreting HEK293 cells (HEK293 WT ORF8) or HEK293 cells containing expression deficient plasmids (HEK293 CO); asterisk indicates WT ORF8 lane; b) Western Blot (WB) of SDS gel in a) and immunoblotting (IB) with an anti-ORF8 antibody (R1).

**Figure S3:**
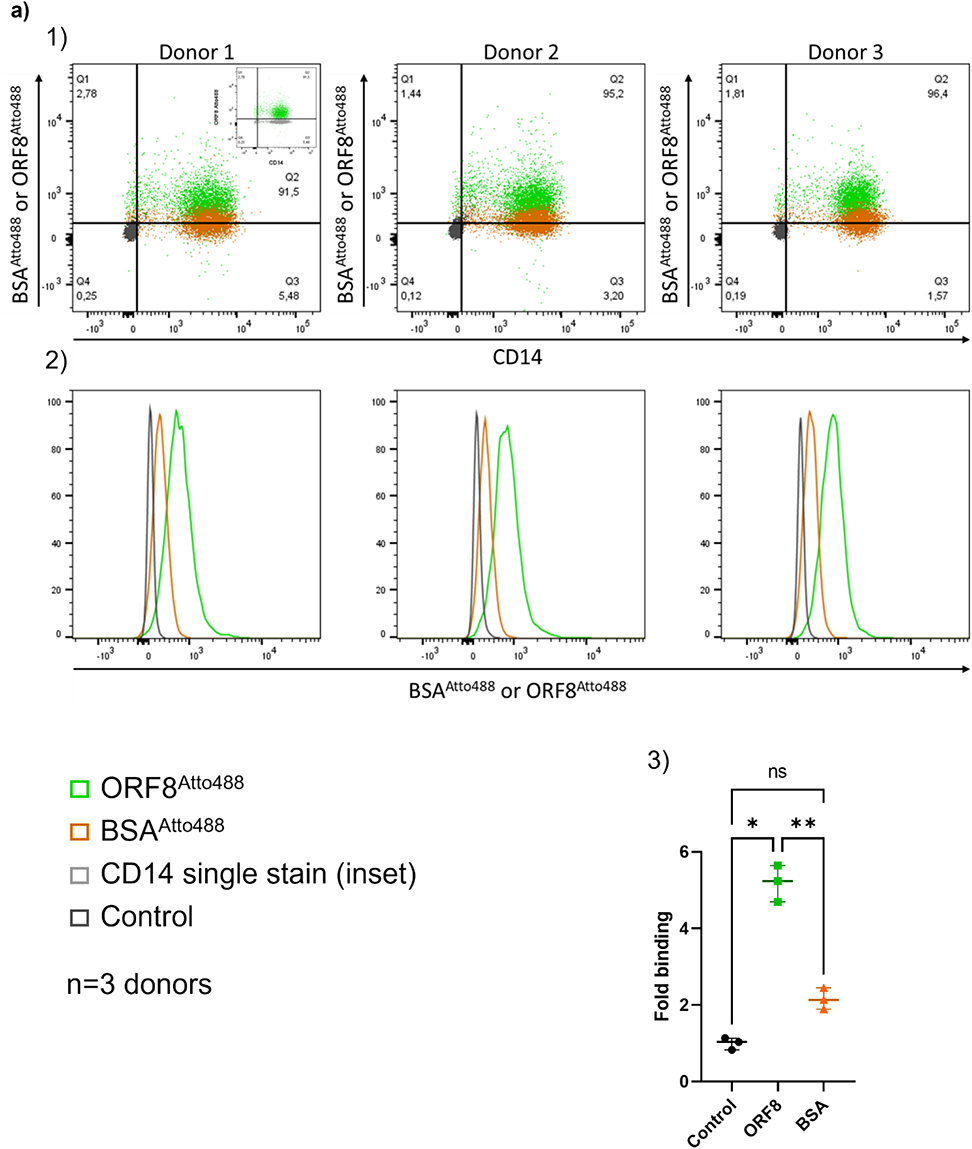
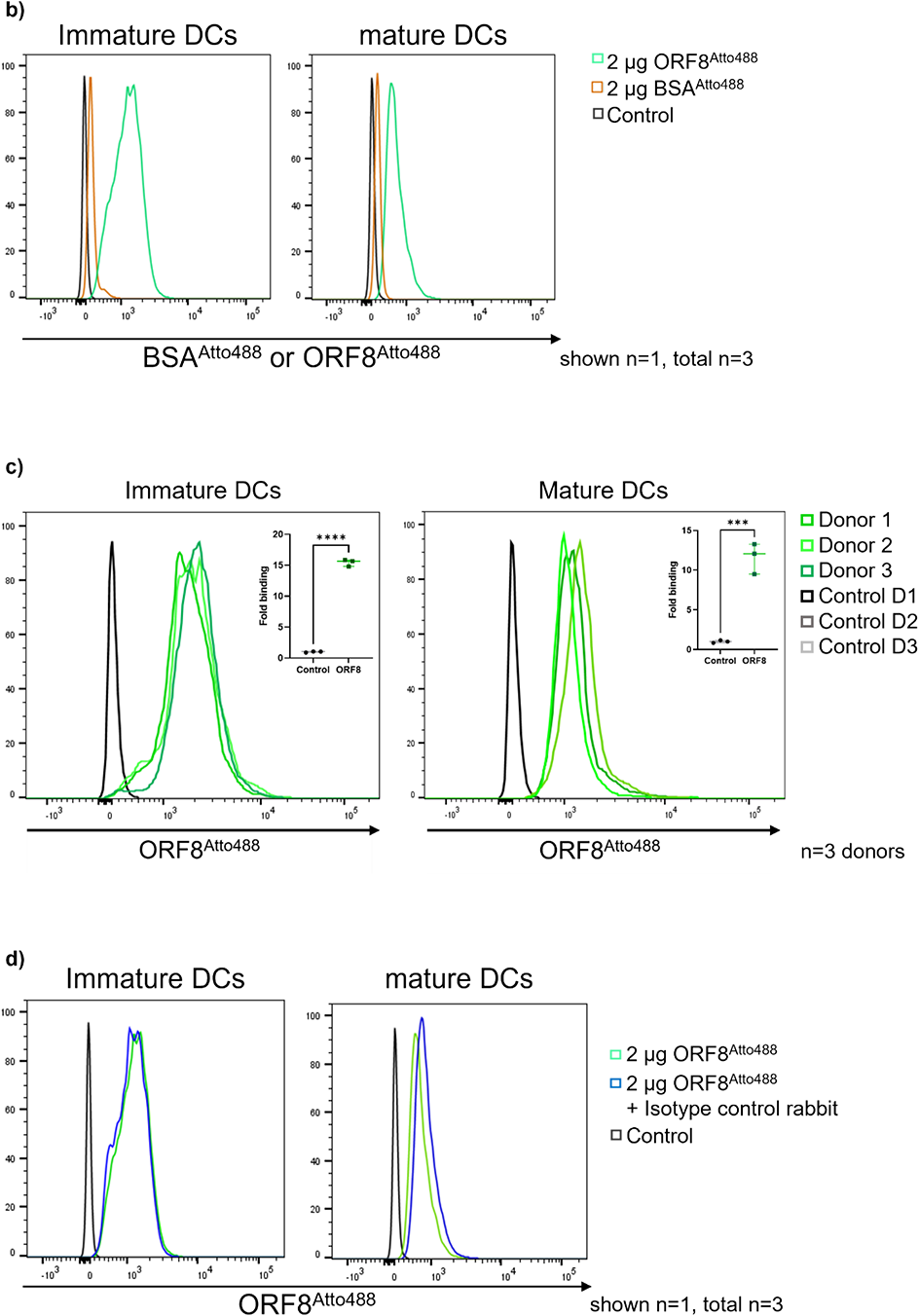
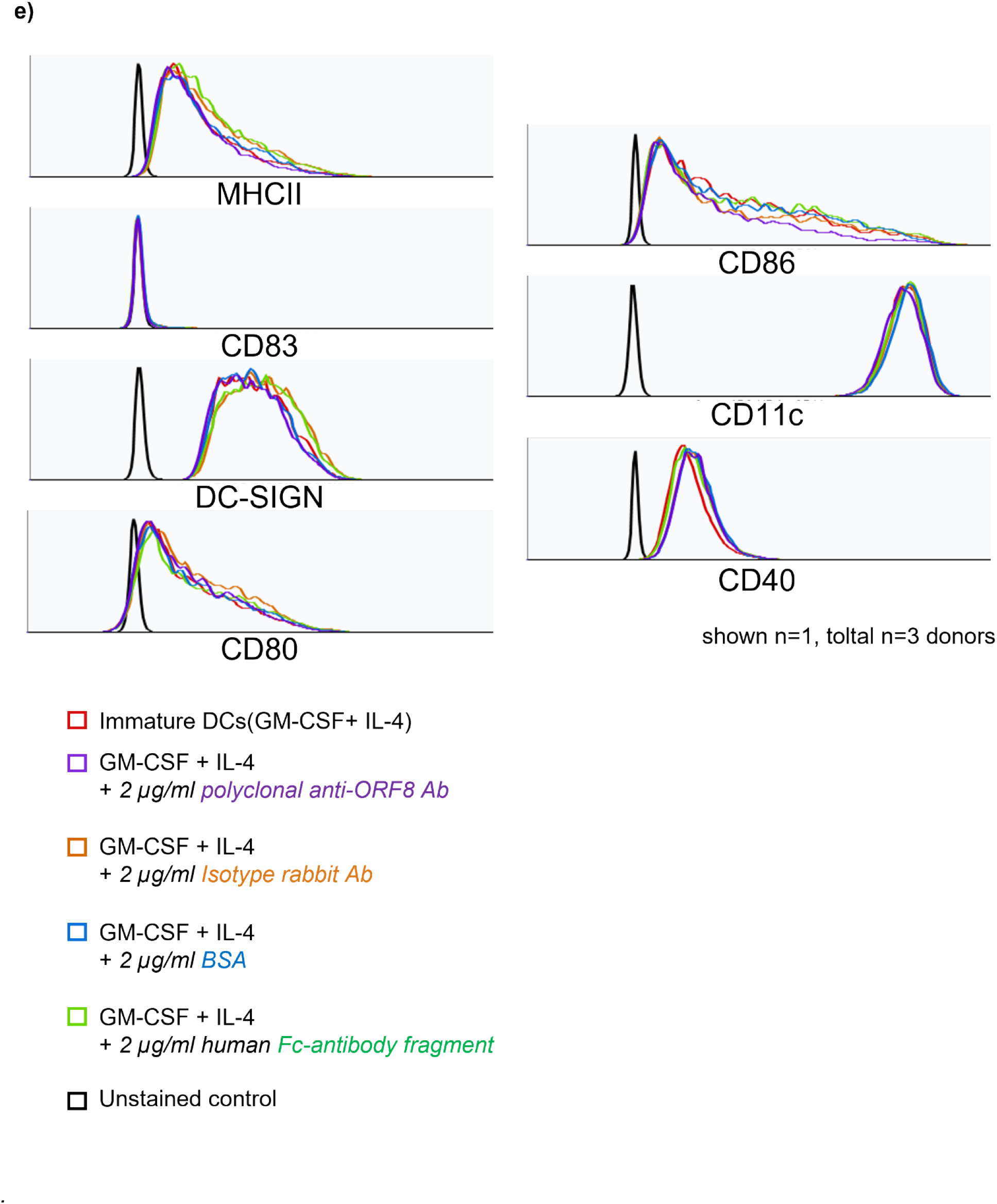
Specific binding of ORF8 to monocytes and dendritic cells. a) Flow cytometric analysis of CD14+ monocytes within PBMCs on the binding of ORF8-Atto488; negative control: unstained cell and BSA-Atto488 as random protein control (1, 2), statistical analysis of the binding (3);. b) The binding capacity of ORF8-Atto488 or BSA-Atto488 to dendritic cells of healthy donors was determined by FACS analysis. c) Binding of ORF8 to immature and mature DCs, shown three individual donors and statistic analysis. d) The effect on ORF8 binding was determined by a rabbit isotype control in FACS analysis. e) Flow cytometry analysis of the effect of control proteins on the expression of MHCII, CD80, CD83, CD86, CD40 and CD11c during differentiation of monocytes into immature dendritic cells with IL-4 and GM-CSF compared to only IL-4+GM-CSF differentiated monocytes. For all experience we performed an analysis of n = 3 independent donors

**Figure S4:**
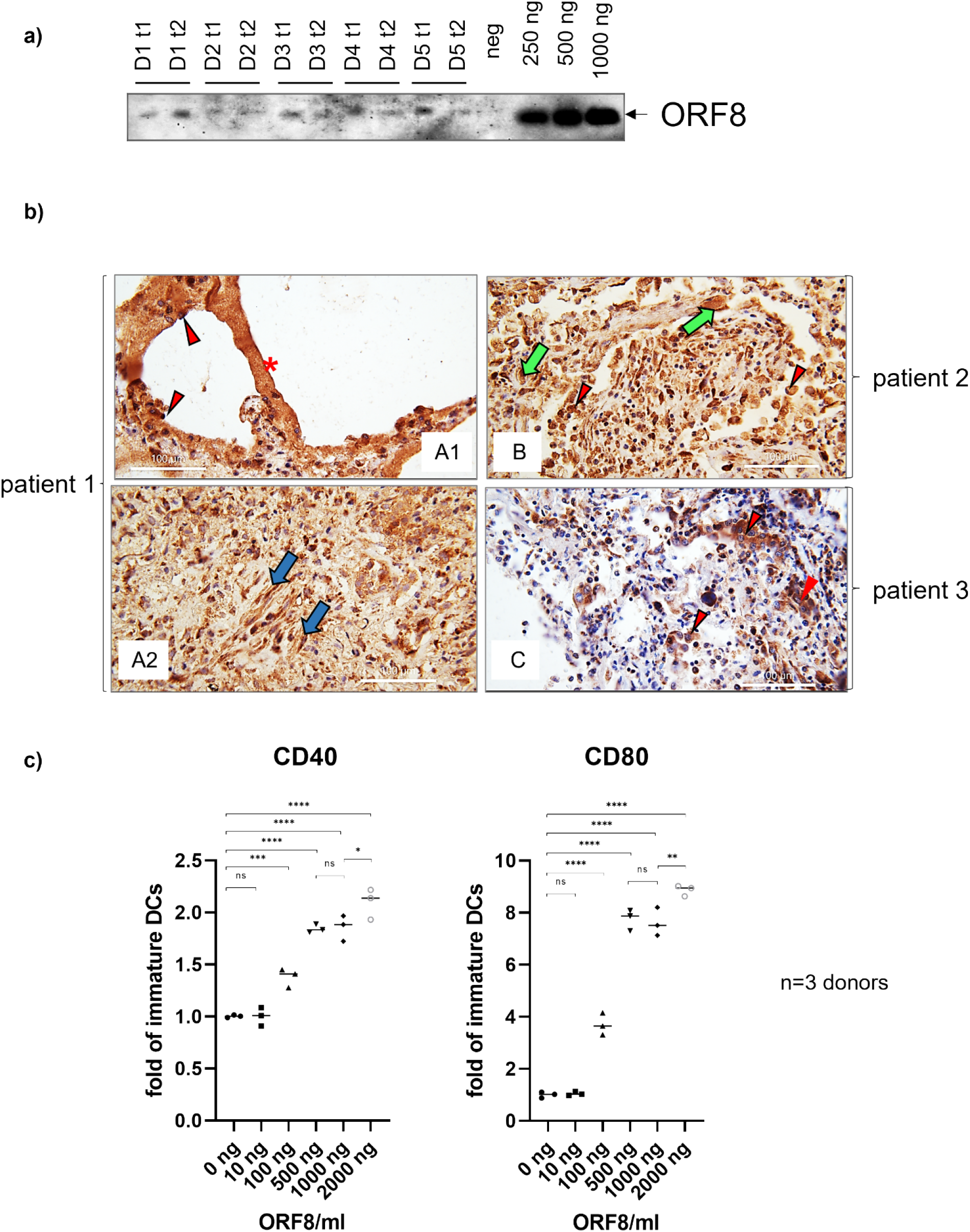
Expression and secretion of ORF8 during infection and dose dependent activation of DCs. a) immuno blot analysis of 2 µl patient sera (n = 5) (two timepoints of hospitalized patients: t1, t2), separated in a 4 – 12 % SDS page. The detection was performed with an rabbit antibody raised against the C-terminus of the ORF8 protein. b) Immunohistochemical detection of ORF8 (a,b,c) and depicts lung changes due to SARS-CoV2 Infection. ORF8 is expressed in hyaline membranes (a1 red*), Pneumocytes (red arrow head) and histiocytes (green arrows) and fibroblastic cells in the fibroblastic foci of the organizing pneumonia (blue arrows) are also positive. Lymphocytes are in all analyzed cases negative, independent of the severity of infection. (n = 3, different severity of infection). c) Monocytes were differentiated to immature dendritic cells by IL-4+GM-CSF in the absence (0 ng /ml) or presence of ORF8 in increasing concentrations (10 – 2000 ng/ml). After 5 days, the expression of maturation marker were analyzed by FACS analysis (n = 3).

**Figure S5:**
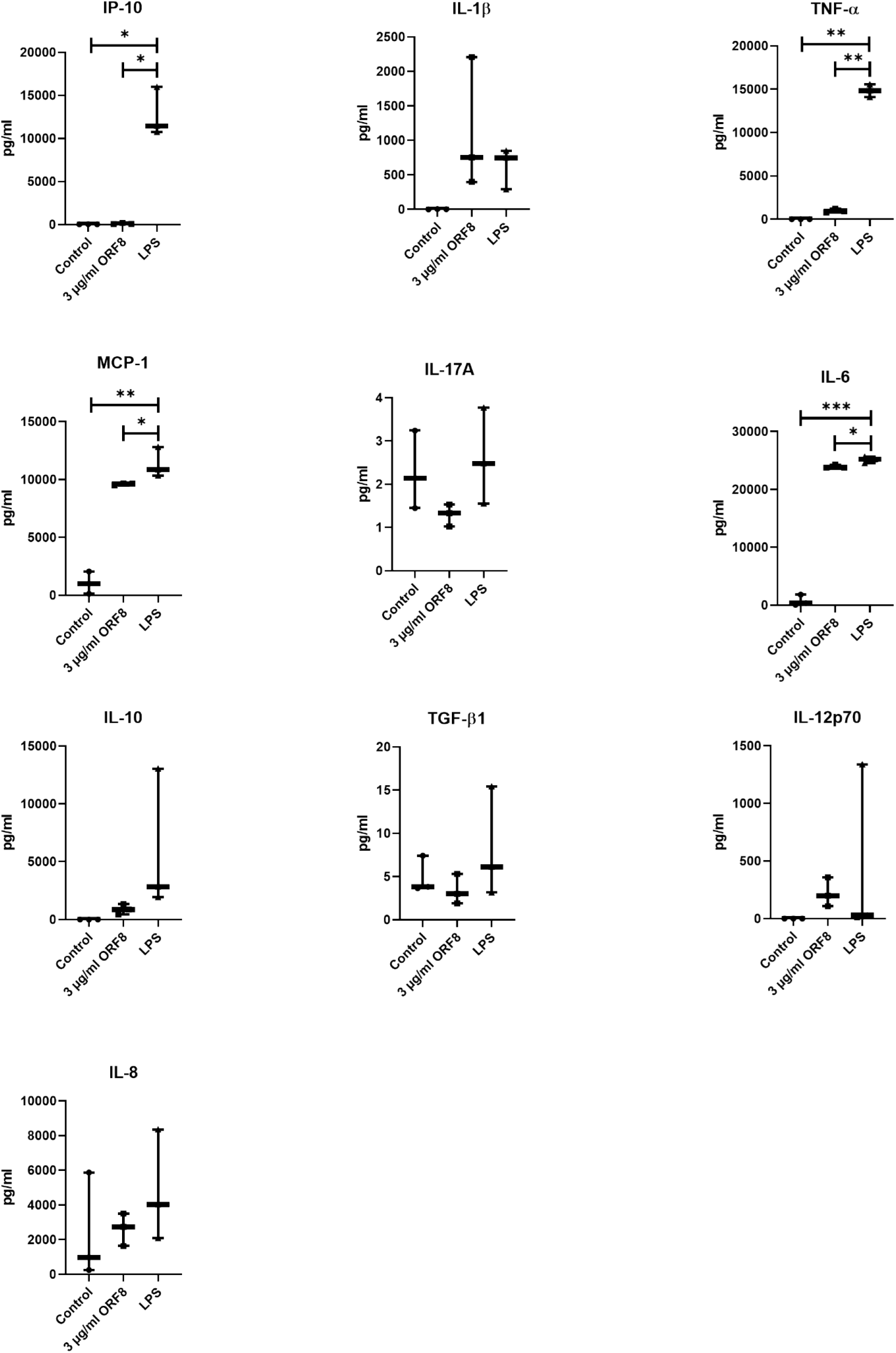
ORF8 shows a unique profile of DC activation that differs from LPS stimulation. Monocytes were differentiated (GM-CSF + IL-4) in the presence or absence of ORF8 (3 µg/ml) or LPS (100 ng/ml). The supernatant of the cells was collected and analyzed with a multiplex assay for the induction of individual chemo-and cytokine profiles of ORF8 respectively LPS in comparison to untreated cells (n=3 of each group).

**Figure S6:**
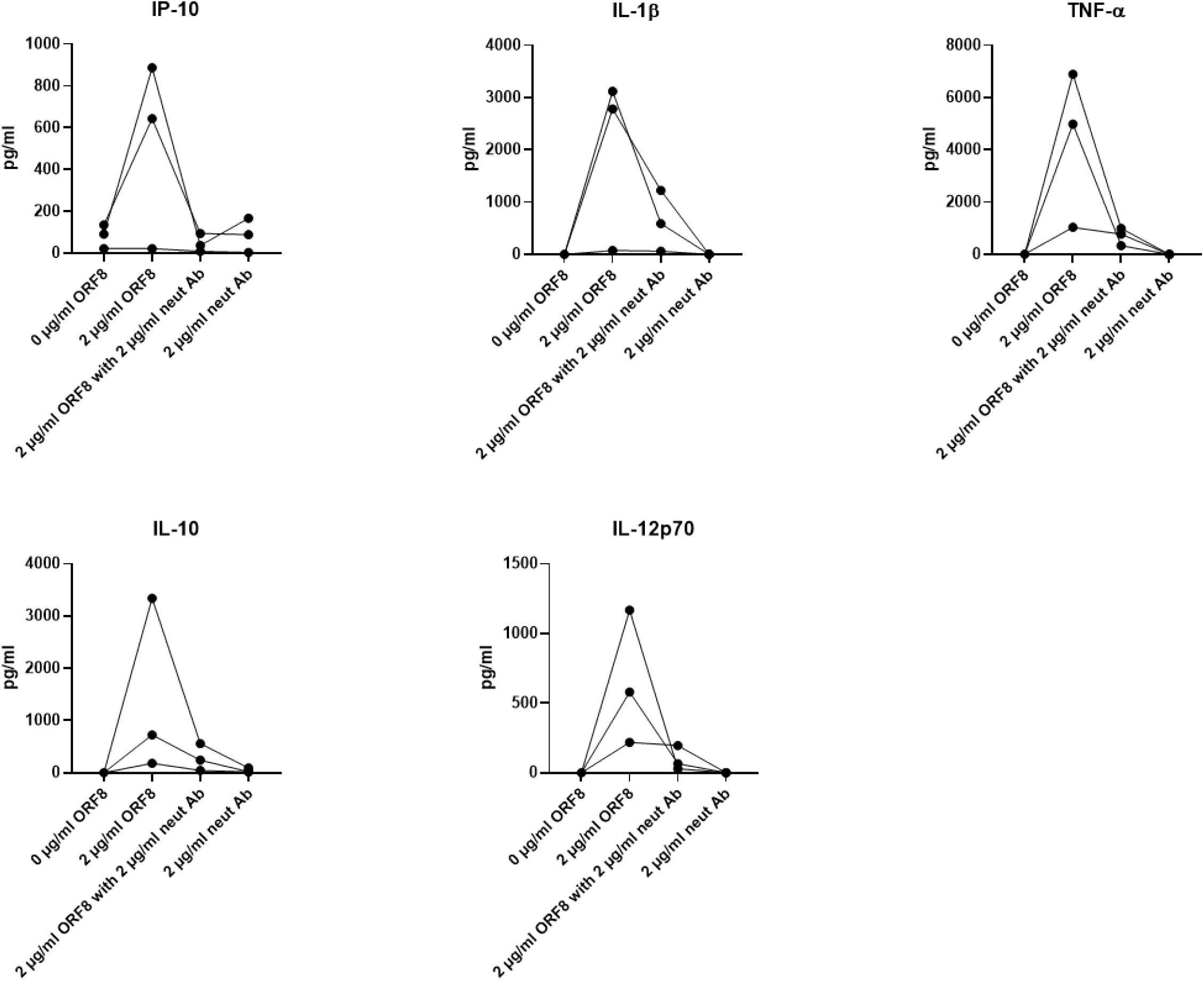
Neutralization of ORF8 blocks the ability of the protein to induce a pro-inflammatory cytokine storm. Monocytes were differentiated (GM-CSF + IL-4) in the presence or absence of ORF8 (2 µg/ml). In addition, the ORF8 protein was neutralized by an anti-ORF8 (2µg/ml) antibody. The supernatant of the cells was collected (n=3 of each group, each line represents one donor) and analyzed with a multiplex assay for the induction of chemo- and cytokines induced by ORF8 and its neutralization (ORF8 is neutralized in the presence of a specific inhibitory antibody).

**Figure S7:**
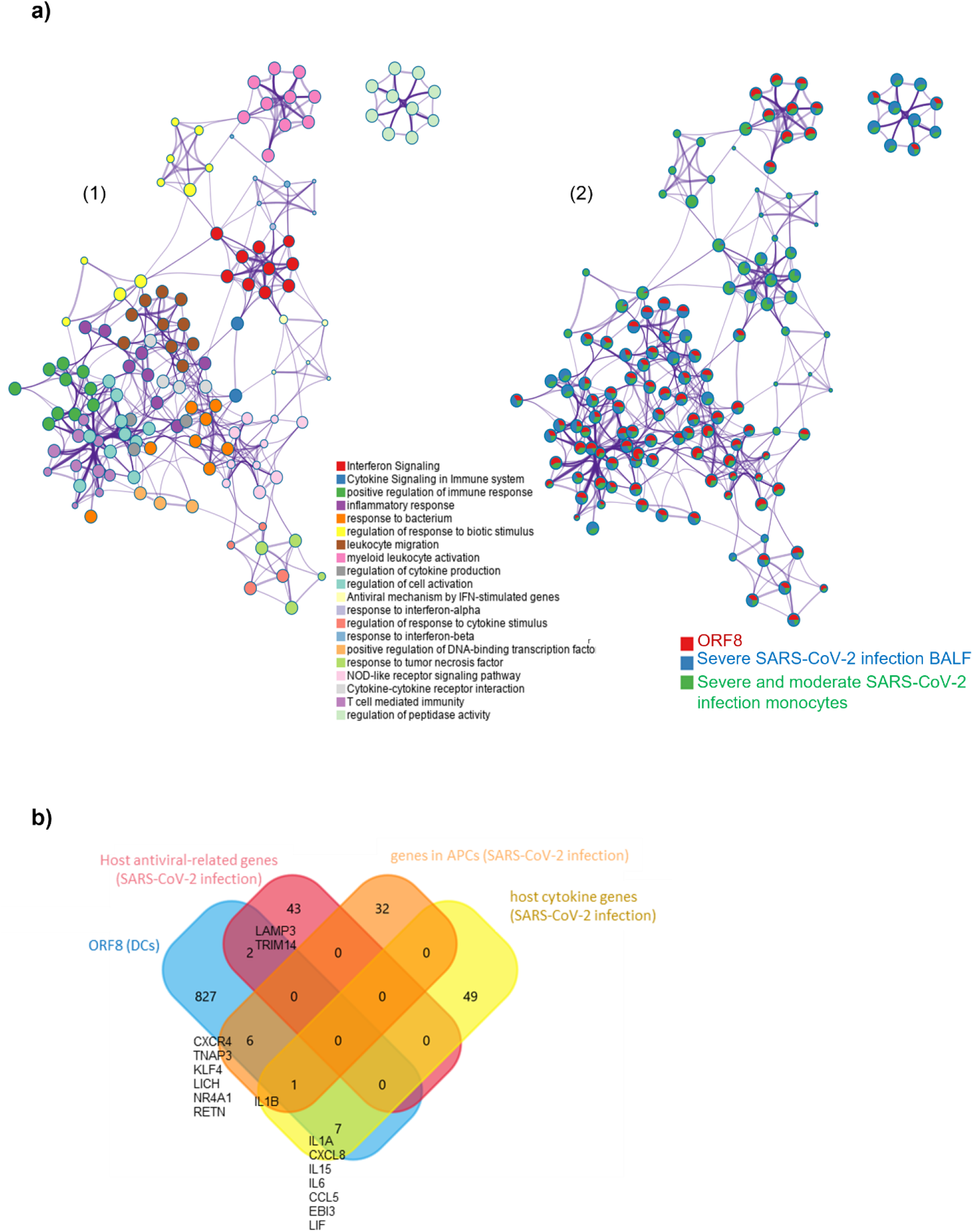
ORF8 induces an inflammatory mRNA profile involved in SARS-CoV-2 infection. a) 1) Network Layout of enriched pathways where each term is represented by a circle node, where its size is proportional to the number of input genes that fall into that term, and its color represents its cluster identity (i.e., nodes of the same color belong to the same cluster). Terms with a similarity score > 0.3 are linked by an edge (the thickness of the edge represents the similarity score). The network is visualized with Cytoscape (v3.1.2) with “force-directed” layout and with edge bundled for clarity. 2) The same enrichment network has its nodes displayed as pies. Each pie sector is proportional to the number of hits originated from the analyzed gene list. The Colour code for the pie sector represents the amount of genes of each list and is consistent with the colors of the figure legend www.metascape.org/COVID. b) Venn plot comparison of detected genes in the ORF8 data set in comparison to published datasets (host antiviral-related genes/host cytokine genes (29); genes in APCs (30)).

**Figure S8:**
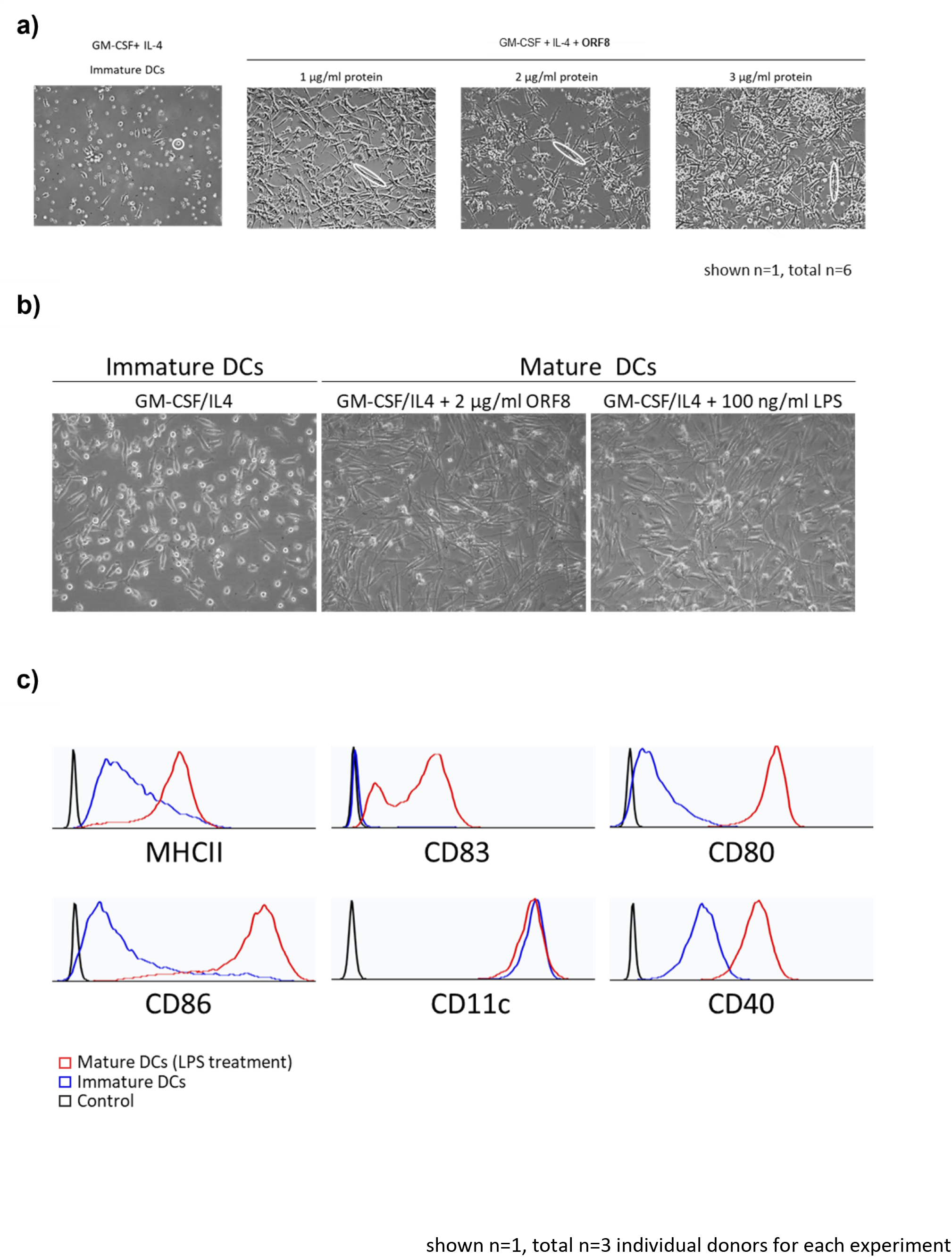
Characterization of immature and mature dendritic cells. a) Effect of ORF8 on the morphology of immature dendritic cells compared to the control group. b) As a positive control for mature dendritic cells, the cells were treated with LPS, an agent that is well characterized to induce maturation of DCs (n=3 of each group, 20x magnification). c) Flow cytometry analysis for MHCII, CD80, CD83, CD86, CD40, and CD11c of immature and mature (LPS treated) DCs.

**Figure S9:**
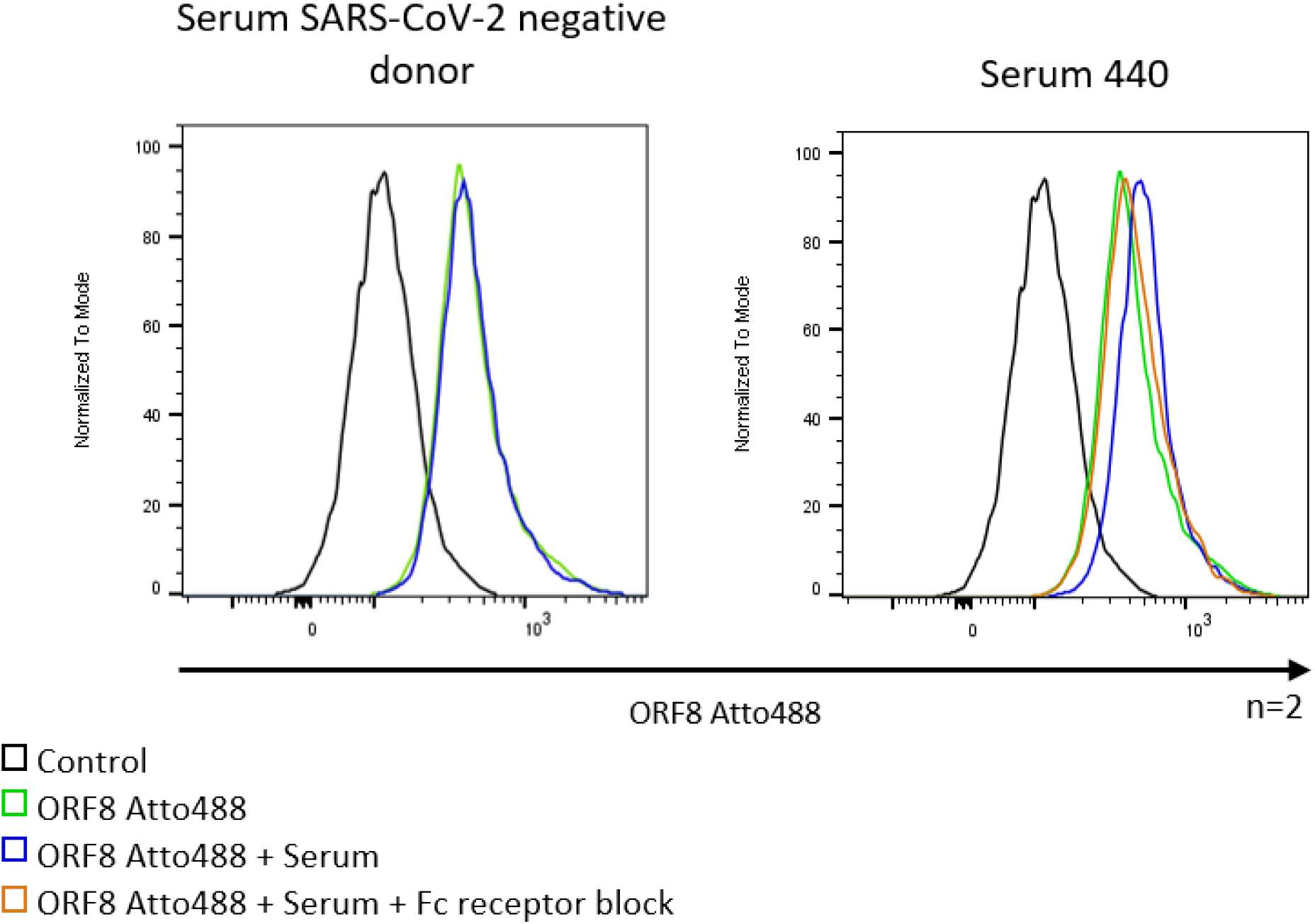
Enhanced binding of ORF8 is antibody-mediated via Fc receptors. The binding capacity of ORF8-Atto488 or ORF8-Atto488 in the presence of sera from anti-ORF8 positive or healthy donors was determined on immature dendritic cells by FACS analysis. The effect of the Fc receptors was determined by blocking them with a commercially available Fc block

**Figure S10:**
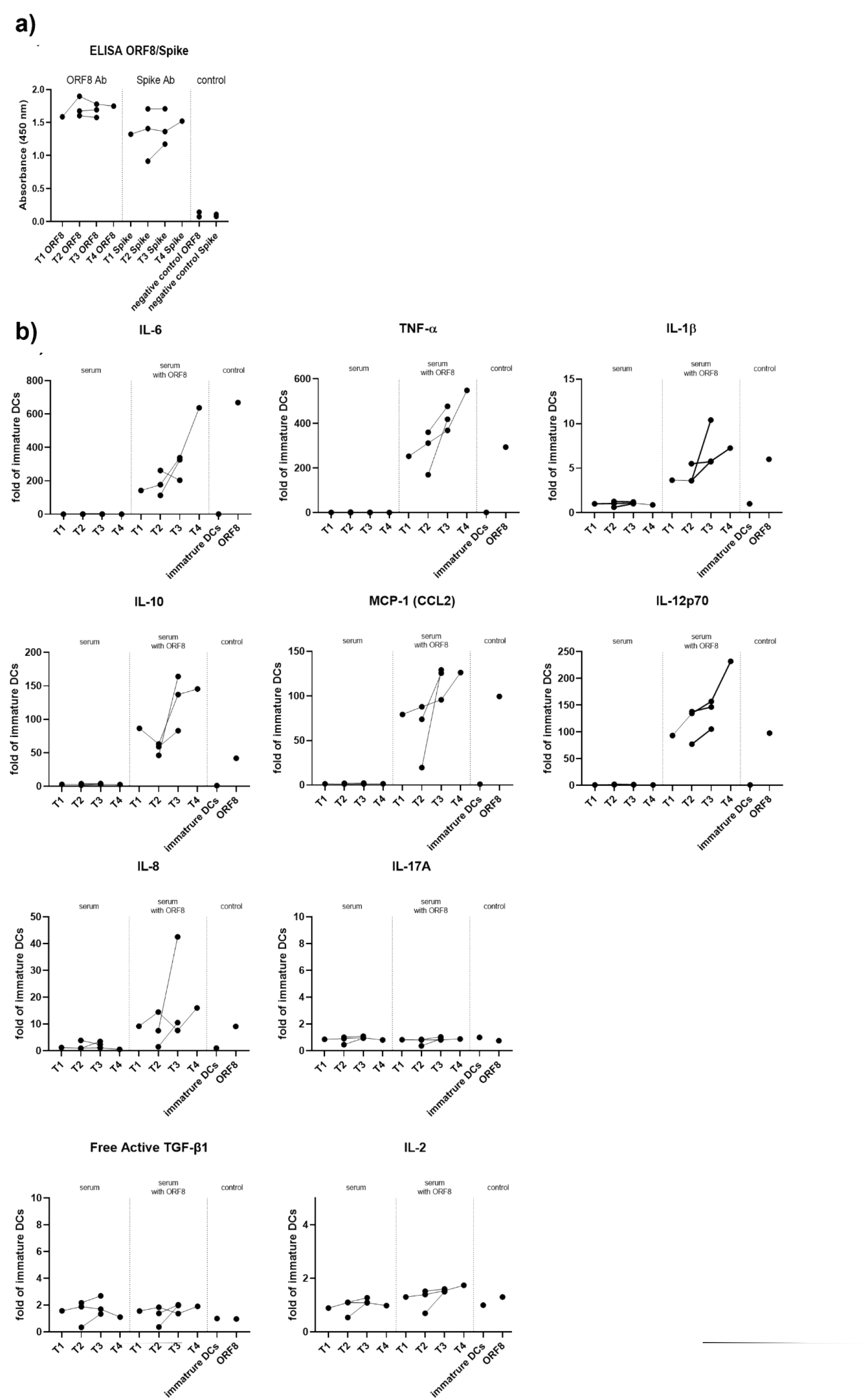
Antibody-dependent enhancement (ADE) of ORF8 is dependent on the time of infection. a) Anti-ORF8 and Spike antibody levels of patients PCR positive for SARS-CoV-2 were detected by an ORF8-ELISA; negative control: plasma of a healthy volunteer (PCR negative tested). b) Monocytes were differentiated (GM-CSF + IL-4) in the presence or absence of ORF8 (2 µg/ml). In addition, the ORF8 protein was treated with patient sera of hospitalized patients from different time points of their infection. As control cells were incubated with sera only. Secreted cytokines and chemokines were analyzed by a multiplex Luminex assay and normalized to fold secretion levels of immature DCs.

**Figure S11:**
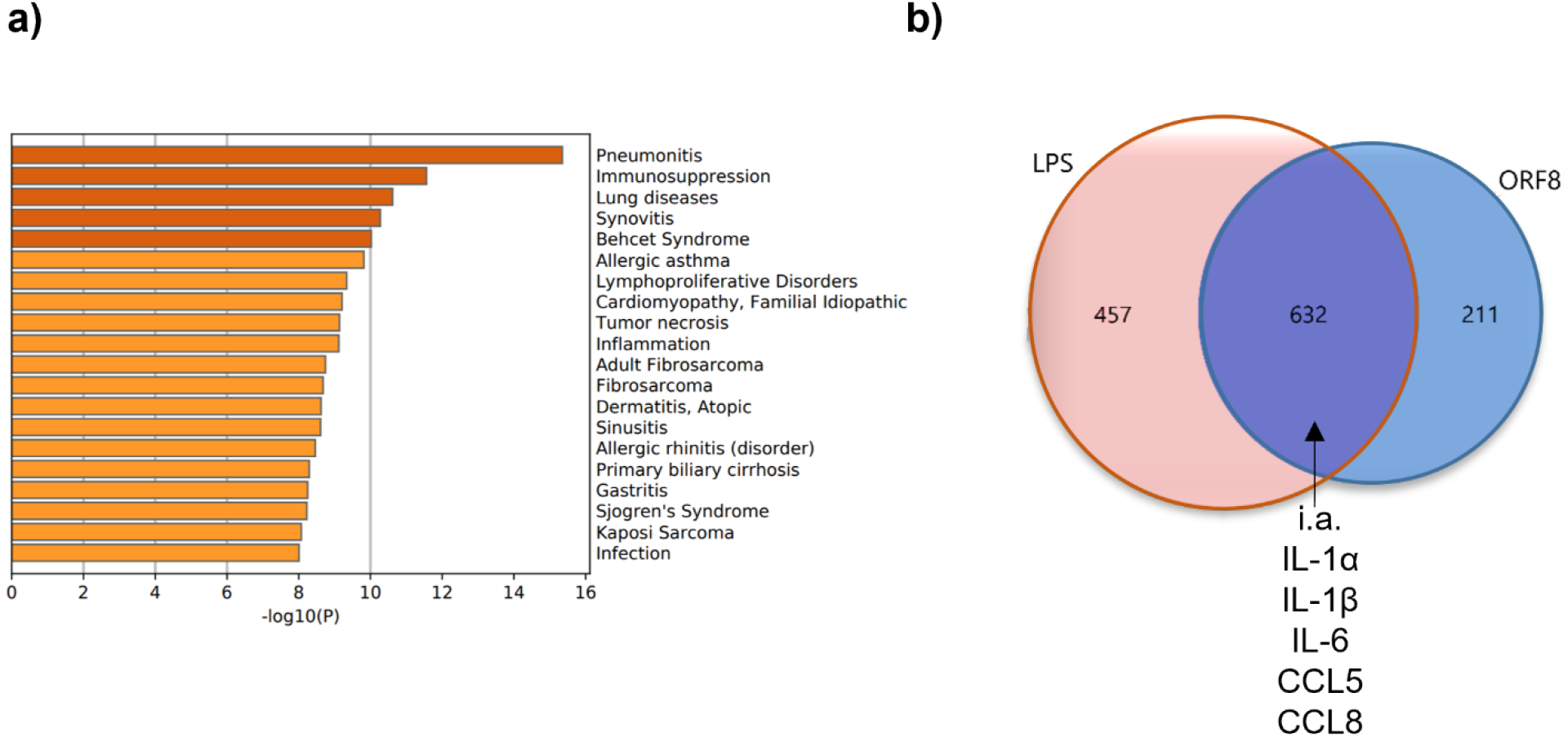
Disease enrichment analysis of ORF8 induced genes and comparison against a well-characterized pro-inflammatory agent (LPS) (endpoint: 120 h). a) Summary of enrichment analysis associated to human diseases using DisGeNET, colored by p-values (https://www.disgenet.org/). b) Van diagram (FunRich) of overlap between ORF8 and LPS (inflammatory stimulus) regulated genes. Overlap between gene lists: purple curves link identical genes and classical inflammatory triggers like IL-6, IL1, CCL8 and CCL5 were marked as found example of an overlapping inflammatory response. 211 regulated RNAs are unique for ORF8.

**Figure S12:**
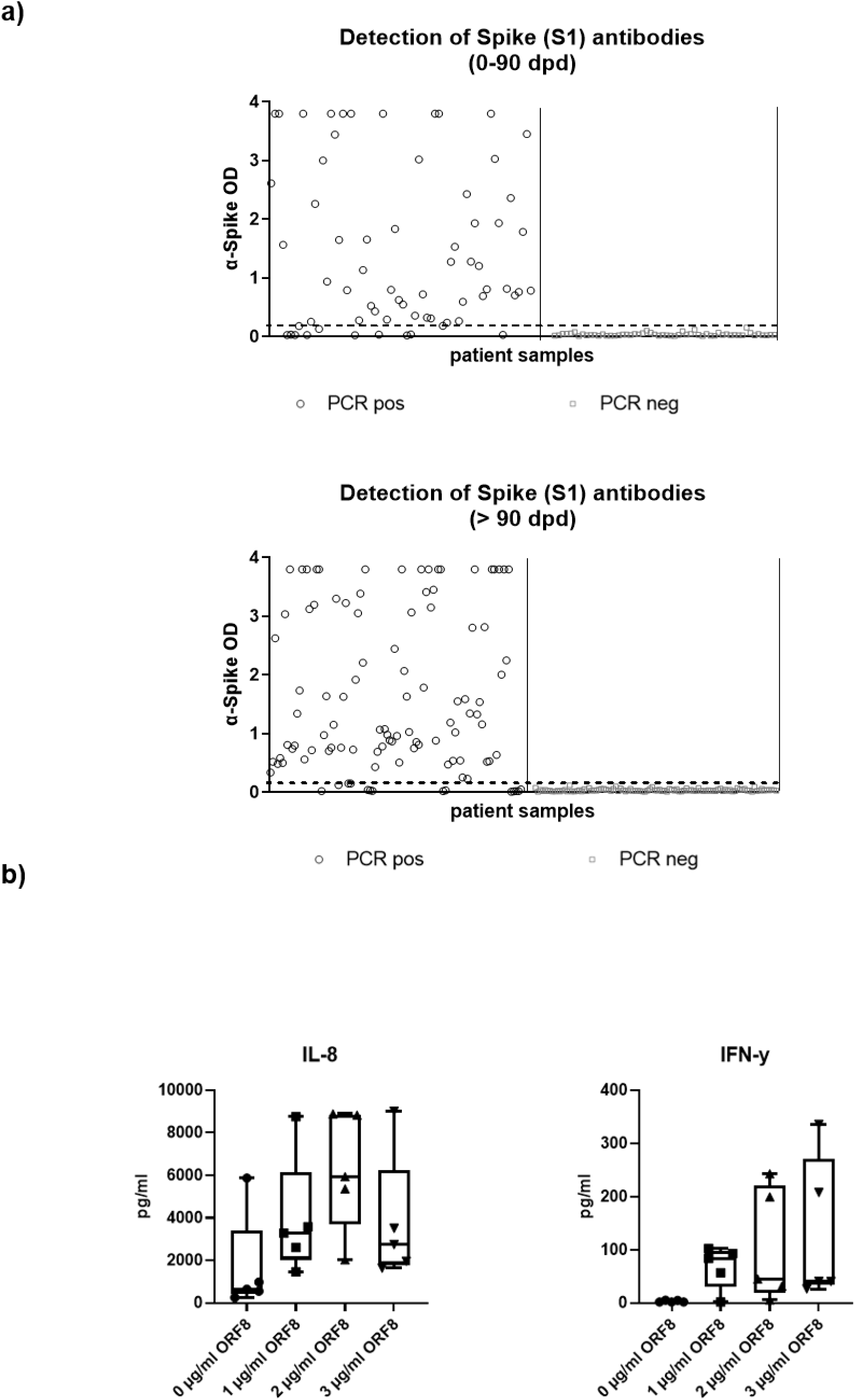
SARS-CoV-2 Spike ELISA. a) ELISA was performed corresponding to the EuroImmun Diagnostic with the corresponding ORF8 patient samples. b) Monocytes were differentiated to immature dendritic cells by IL-4 + GM-CSF in the absence (0 ng /ml) or presence of ORF8 in increasing concentrations (1 – 3 µg/ml). After 5 days, the supernatant was collected, and cytokine and chemokine levels were determined using a cytokine bead array (n = 5). IL-8, Interleukin 8; IFNγ, interferon γ

**Figure S13:**
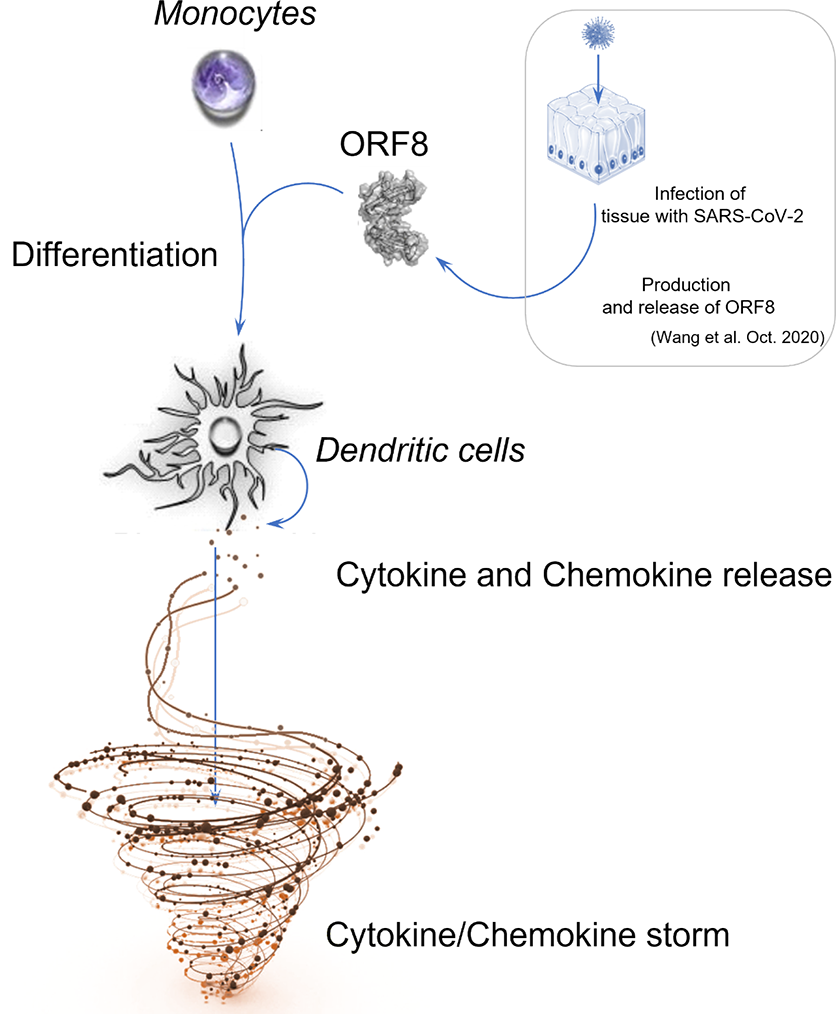
Graphical Abstract.

## Notes

### Competing Interest Statement

The authors have declared no competing interest.

